# A Leadfield-Free Optimization Framework for Transcranially Applied Electric Currents

**DOI:** 10.1101/2024.12.18.629095

**Authors:** Konstantin Weise, Kristoffer H. Madsen, Torge Worbs, Thomas R. Knösche, Anders Korshøj, Axel Thielscher

**Affiliations:** Department of Clinical Medicine, Aarhus University, Aarhus, Denmark; Methods and Development Group “Brain Networks”, Max Planck Institute for Human Cognitive and Brain Sciences, Leipzig, Germany; Leipzig University of Applied Sciences (HTWK), Institute for Electrical Power Engineering, Leipzig, Germany; Technical University of Denmark, Section for Cognitive Systems, Department of Applied Mathematics and Computer Science, Kongens Lyngby, Denmark; Danish Research Centre for Magnetic Resonance, Department of Radiology and Nuclear Medicine, Copenhagen University Hospital Amager and Hvidovre, Hvidovre, Denmark; Department of Neurosurgery, Aarhus University Hospital, Aarhus, Denmark; Technical University of Denmark, Section for Magnetic Resonance, Department of Health Technology, Kongens Lyngby, Denmark

**Keywords:** Electroconvulsive Therapy, Montage Optimization, Temporal Interference Stimulation, Transcranial Electric Stimulation, Tumor Treating Fields

## Abstract

**Background:** Transcranial Electrical Stimulation (TES), Temporal Interference Stimulation (TIS), Electroconvulsive Therapy (ECT) and Tumor Treating Fields (TTFields) are based on the application of electric current patterns to the brain.

**Objective:** The optimal electrode positions, shapes and alignments for generating a desired current pattern in the brain vary between persons due to anatomical variability. The aim is to develop a flexible and efficient computational approach to determine individually optimal montages based on electric field simulations.

**Methods:** We propose a leadfield-free optimization framework that allows the electrodes to be placed freely on the head surface. It is designed for the optimization of montages with a low to moderate number of spatially extended electrodes or electrode arrays. Spatial overlaps are systematically prevented during optimization, enabling arbitrary electrode shapes and configurations. The approach supports maximizing the field intensity in target region-of-interests (ROI) and optimizing for a desired focality-intensity tradeoff.

**Results:** We demonstrate montage optimization for standard two-electrode TES, focal center-surround TES, TIS, ECT and TTFields. Comparisons against reference simulations are used to validate the performance of the algorithm. The system requirements are kept moderate, allowing the optimization to run on regular notebooks and promoting its use in basic and clinical research.

**Conclusion(s):** The new framework complements existing optimization methods that require small electrodes, a predetermined discretization of the electrode positions on the scalp and work best for multi-channel systems. It strongly extends the possibilities to optimize electrode montages towards application-specific aims and supports researchers in discovering innovative stimulation schemes. The framework is available in SimNIBS.

## 1. Introduction

Transcranial electric stimulation (TES), temporal interference stimulation (TIS), electroconvulsive therapy (ECT), and tumor treating fields (TTFields) are therapeutic modalities that deliver electric currents to the brain by means of electrodes attached to the scalp and have been investigated for a range of neurological and oncological conditions (Bestmann et al., 2017; Mun et al. 2018; Chakrabarti et al., 2010; Wenger et al., 2018). Despite their differences in stimulation parameters and intended applications, they share the need for a spatially precise delivery of electric currents to specific regions of the brain or tumor tissue.

State-of-the-art algorithms for the optimization of the electric current patterns applied to the brain are based on leadfields (Dmochowski et al., 2017; Saturnino et al., 2019; Saturnino et al., 2021). They require a predetermined discretization of the potential electrode positions on the scalp, similar to an EEG cap, and small electrodes to prevent spatial overlaps between neighboring positions. A dense discretization and multiple stimulation channels are required to reach the best possible spatial resolution (Saturnino et al., 2019). This comes with potential drawbacks, such as increased costs, complexity and setup time, which can reduce their benefits in practice. Thus, despite the theoretical advantages of multi-channel systems, systems with a low number of channels are often used in practice. However, a systematic framework to determine the optimal electrode shapes and positions for standard TES applications and TTFields has been lacking so far. Leadfield-based approaches are not well suited in these cases, because spatially extended electrodes and complex electrode arrays cannot be easily mapped to the discrete positions of the cap layout. Instead, an optimization routine is required that is capable of “moving” electrodes freely over the head surface while avoiding mutual intersections in order to determine the optimal electrode positions, orientations and alignments.

To achieve this, we developed a flexible optimization framework that allows for a fast and computationally efficient determination of optimal electrode configurations considering individual head and cortical anatomies. It builds upon a geodesic coordinate system for the representation of electrode configurations on the scalp surface, resulting in compact descriptions of the optimization problem with a low to moderate number of parameters. In addition, it avoids the need to explicitly model the electrodes by approximating the current patterns injected at the electrode-skin interfaces (Miranda et al., 2006; Korshoej et al., 2018), allowing for an efficient forward modeling of the electric fields based on the Finite-Element Method. It enables the maximization of the intensity in a target area of the brain or in tumor tissue. Alternatively, the optimization of the intensity-focality trade-off (Fernandez-Corazza et al., 2020) to reach a desired balance between the target intensity and the spread to other areas is supported in a computationally efficiently way using receiver operating characteristic (ROC) curves. In combination, these advances result in optimization problems that can be successfully tackled with established solvers on standard computer hardware.

We validate the performance of the optimization framework through a series of different application scenarios, starting with standard unfocal TES with two rectangular electrodes as well as focal TES with center-surround montages (4×1 TES). For TIS, the locations of two electrode pairs operating at different frequencies are optimized to target deep brain structures by maximizing the envelope of the low-frequency interference pattern in the target region. Moreover, for TTFields, the electric field in tumor tissue is maximized to inhibit tumor growth. Finally, we briefly showcase the potential of the method in two further application scenarios, namely the optimization of the electrode geometries of standard two-electrode TES, and the optimization of ECT compared to a standard right unilateral (RUL) montage (Martin et al., 2021).

The new method complements lead-field-based approaches and allows clinicians to personalize stimulation in a range of clinically tested applications, potentially improving therapeutic efficacy while minimizing the risk of adverse effects. It can be integrated into the daily routine of laboratories and clinics using standard computer hardware. It does, however, benefit from a neuronavigation system, because the electrode positions and orientations are free and not restricted to an EEG 10-20 system.

## 2. Methods

### 2.1. Overview of the Optimization Framework

Our approach directly optimizes the relevant parameters, such as the center positions and orientations of rectangular electrodes for standard unfocal TES, to maximize the desired goal function. The latter could be the average electric field strength in a target brain region or the focality of the electric field in case of TES, but also more complex functions such as the electric field envelope, averaged in a target region for TIS. Further, the optimization is subject to a number of necessary constraints, such as avoiding positions that overlap with the face area.

Numerical optimization requires the repeated evaluation of the goal function for varying parameter combinations, which is in practice only feasible when the computational costs per evaluation are low. So far, this has prevented the direct use of electric field calculations as part of optimization procedures for TES, TIS and TTFields. For example, a SimNIBS FEM (Finite Element Method) simulation includes electrode modeling, electric field calculations and post processing, which can take minutes, depending on the geometry of the head model and the applied electrodes, prohibiting fast iterative updates. Our new framework overcomes this problem by combining the following approaches:

1. The electrodes are approximated by current patterns that are injected directly into the skin surface. This avoids the need for adding the electrode models to the head mesh. In addition, as the head mesh stays unchanged, costly preparation steps for the FEM calculations can be reused to speed up repeated simulations when varying the parameters.
2. In particular larger electrodes have a non-uniform current density at the electrode-skin interface, with higher current densities towards the electrode edges (Miranda et al., 2006; Korshoej et al. 2018). Here, the injected current patterns are dynamically weighted to approximate this effect and maintain simulation accuracy.
3. A suitable ellipsoidal coordinate system is created by fitting a tri-axial ellipsoid to the individual upper head shape in order to support an efficient parametrization of the search space. Pre-calculation of the mapping between the coordinate system and the skin positions makes it computationally efficient to determine the current injection pattern on the skin for a new parameter set during optimization.

In combination, these steps reduce the time per evaluation of the goal function to less than one second on a standard modern computer for our implementation. The following sections outline the details of the optimization framework. Additional in-depth information is given in the Supplementary Material S1 and S2.

### 2.2. Electric Field Calculations

We base our approach on the first-order tetrahedral Finite Element method with so-called super-convergent patch recovery (SPR) implemented in SimNIBS (Saturnino et al., 2019) that has been validated to have a good tradeoff between computational costs and accuracy. In FEM-based TES simulations, electrodes are usually modeled as additional volumes, which are merged with the head model, followed by an adaptation of the tetrahedral mesh to the new geometry. Changing the mesh additionally requires that the preparation steps for the FEM (i.e., the creation and pre-conditioning of the FEM stiffness matrix [**A**]) are repeated as well. Overall, this can take several minutes in the current SimNIBS implementation. In general, these limitations are not restricted to this specific FEM implementation but apply to varying extent also to alternative methods such as the Finite Difference Method or the Boundary Element Methods.

Here, we avoid repeated costly preparation steps and approximate the electrodes by defining current sources directly at the electrode-skin interface (*I_1_*, *I_2_*,…in Fig. 1), using von Neumann boundary conditions at the corresponding skin nodes. This reduces the calculations required for an update of the FEM source vector **b** to bring it in line with the new boundary conditions, followed by solving the updated matrix equation [**A**]**v = b** to get the new solution for the electric potential **v.** The goal function value is then calculated from the FEM solution for **v** by evaluating the electric field or a derived measure in one or more regions of interest (ROI). This involves taking the numerical gradient of the **v** in the tetrahedra intersecting with the ROI, followed by interpolating the field to the points that comprise the ROI. The interpolation uses superconvergent patch recovery (SPR) (Zienkiewicz and Zhu, 1992) and is implemented computationally efficiently as multiplication with a pre-calculated sparse weight matrix, similar to the approach outlined in Cao, Madsen et al. (2024) for TMS. In combination, these optimizations make evaluations of the cost function for new parameter sets possible within one second or less. In our implementation, ROIs can be defined as scattered point clouds, by the nodes of triangulated surfaces, or by sub-volumes of the tetrahedral mesh. To-be-avoided brain areas can be similarly specified by defining non-ROIs, which is required, e.g. when aiming to optimize the focality of the injected electric field.

**Fig. 1:**
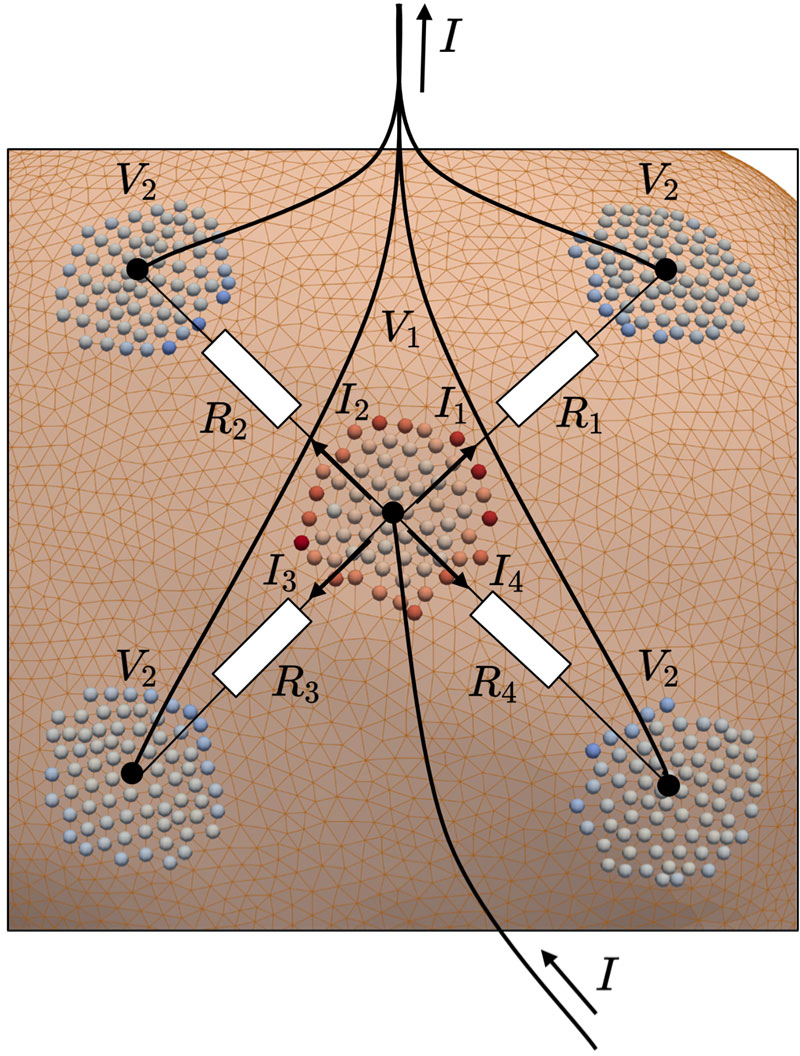
Example of the definition of nodal current sources and electrical equivalent circuit according to an exemplary focal 4×1 TES montage. The total current *I* is injected in the inner electrode and distributes over the outer electrodes (*I*_l_… *I*_4_). The underlying head and brain anatomy defines the equivalent resistances (*R*_l_…*R*_4_). If the outer electrodes are connected to a single channel, the voltages over the resistances (*ΔV* = *V*_l_ - *V*_2_) are equal.

While the above approach is very efficient, it requires setting the node currents *I_1_, I_2_, etc.* correctly to account for the non-uniform current density distribution that occurs at the interface area and maintain the simulation accuracy also in case of larger electrodes. In addition, when several electrodes are connected to the same stimulation channel (as in case of TTFields), the currents at the skin interfaces of those electrodes influence each other. Our solution to maintain accurate estimates of the node currents also in those cases is described in Supplementary Material S1. We refer to it as *node-wise or electrode-wise Dirichlet correction,* depending on whether it is applied to adjust all node currents individually, or just ensures the correct amount of current for each electrode connected to a common channel (the latter is computationally less demanding).

For setting current sources at the electrode-skin interface, our approach involves an efficient identification of the skin surface nodes that correspond to the given electrode positions, orientations and shapes. During preparation, a tri-axial ellipsoid is fitted to the skin region in the upper part of the head (Fig. 2b) and the mapping between positions on the ellipsoid and the skin surface is established by projecting rays from the ellipsoid along its normal direction towards the head surface. This way, determining the skin position corresponding to a given spherical coordinate (θ’, φ′) on the ellipsoid is straightforward. During optimization, the center positions of the electrodes (or electrode arrays) are parameterized as spherical coordinates and their orientation by the angle α′ relative to the vector of constant φ in the ellipsoidal space. This approach enables the straightforward definition of electrode shapes and array layouts in a two-dimensional planar coordinate system (Fig. 2a). During optimization, the shapes and layouts can be efficiently mapped to the ellipsoid by solving the associated direct geodesic problem, known from differential geometry. In combination, this makes the computation time for determining the surface nodes in each iteration of the optimization comparatively short, with around 0.05 sec for a 4×1 TES montage with 4 external electrodes. Details of the geodesic coordinate system are given in Supplementary Material S2 and the stability of the fitting procedure for different head shapes is validated in Supplementary Material S4.

**Fig. 2:**
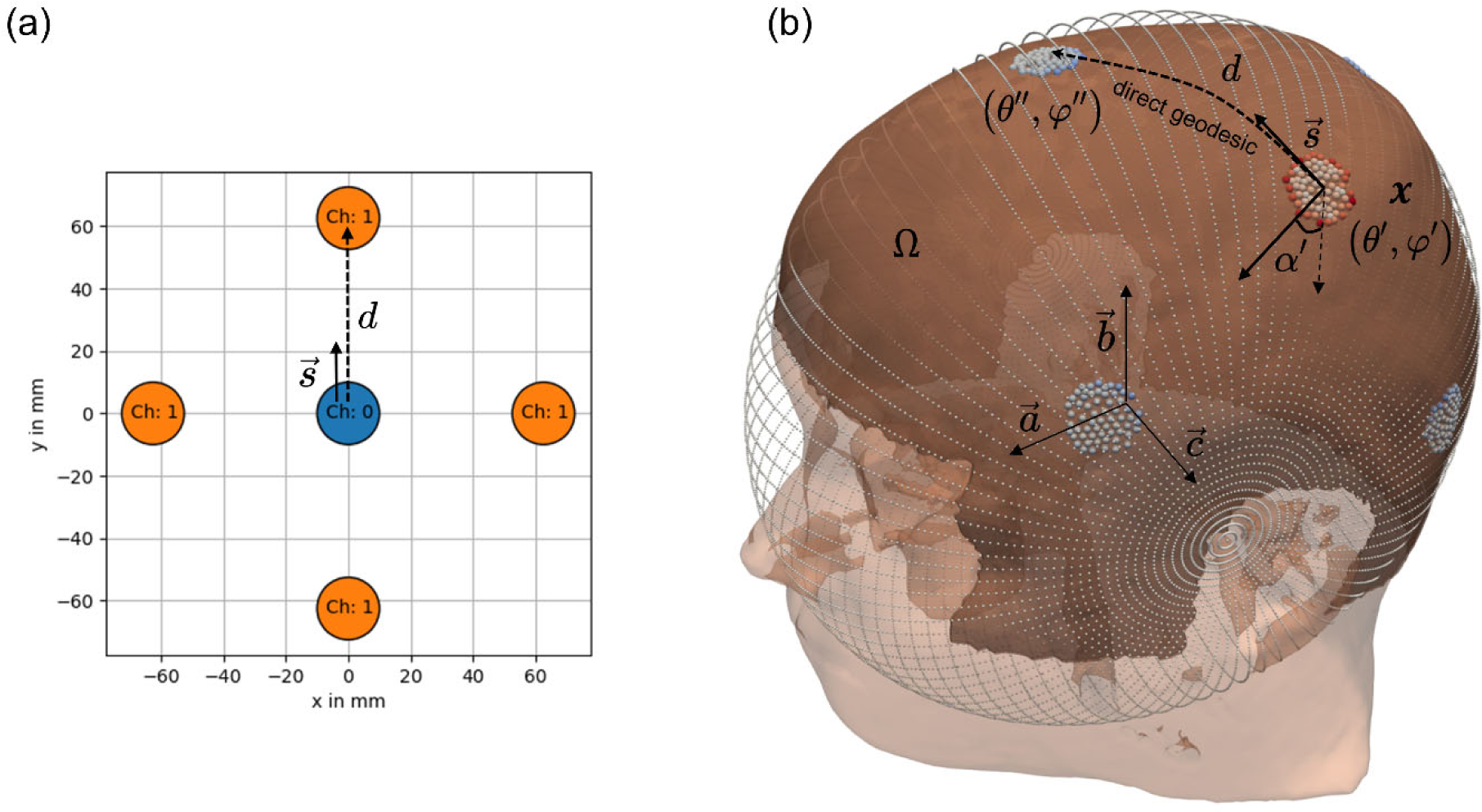
(a) Base geometry of the electrode array defined by the user in normalized space (*xy*-plane); (b) Head model showing the valid region on the skin surface *Ω*, where the electrodes can be located (darker region). The fitted triaxial ellipsoid used to parametrize the electrode array location and orientation (*x*) for the optimization is indicated with small gray dots. In- and output currents are defined in the skin nodes (red and blue dots, respectively) according to the applied Dirichlet approximation to consider the equal voltage constraint of each electrode channel.

### 2.3. Goal Functions

The definitions of the goal functions are based on the magnitude (i.e. strength) of the electric field |**E**| in the specified ROIs (and non-ROIs). Alternatively, the electric field component *E_n_* that is locally orthogonal to the cortical sheet (given by the normal vector **n**) can be used to optimize the in- or outwards pointing field component. For TIS, two stimulation channels are active simultaneously and create a superposition of two electric fields, **E**^(1)^ and **E**^(2)^, oscillating at slightly different frequencies. In this case, the goal function is based on the maximal amplitude of the modulation envelope of the superimposed fields (Grossmann et al., 2017):

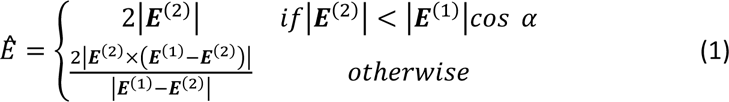

Alternatively, the maximal amplitude of the modulation envelope along a specific direction of interest, indicated by unit vector ***u*** can be used (Grossmann et al., 2017):

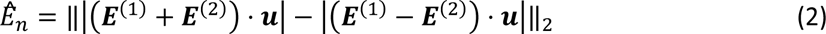

In the following definitions of the implemented goal functions, the field magnitude |**E**|, its normal component *E_n_*, and the TIS envelope magnitudes ^Ê^ and *Ê_n_* can be used interchangeably, depending on the intended application and optimization goal, and are therefore commonly denoted as *target measure E*.

#### 2.3.1. Intensity-based goal functions

For maximizing the intensity of the target measure in the ROI, we use the negative of its average in the ROI as goal function during the minimization process:

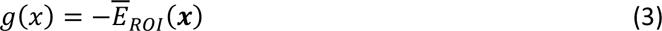

Vector ***x*** denotes the to-be-optimized parameters such as the electrode centers.

For TTFields, two pairs of electrode arrays are used, which are switched on and off alternately. Applying the simple assumption that the combined treatment effect is the average of the effects of the two electric fields, the goal functions are calculated individually for each stimulation pair and their average value is used (Korshoej et al., 2018):

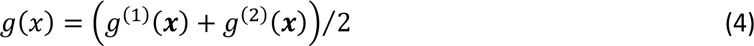

#### 2.3.2. Focality-based goal functions

The aim of optimizing the focality is usually to strengthen the target measure in the ROI, while at the same time reducing it in non-ROI areas. Prior studies on multi-channel TES demonstrated that, when constraining the total injected current to ensure safety, maximizing the target measure in the ROI coincides with a reduction of focality (Fernandez-Corazza et al., 2020). The relationship was approximately sigmoidal, i.e. slight further increases of already high intensities were accompanied with large decreases in focality. Formally, this represents a Pareto front between maximum intensity and maximum focality, whereby it is difficult for the user to select the hyperparameters of the optimizer in order to achieve an individually optimal result.

Here, we use the receiving operating characteristic (ROC) curve to evaluate the intensity-focality tradeoff (Fig. 3a). For that, the target measure in the ROI and in the non-ROI are compared to user-defined thresholds and the relative number of elements that fulfill the conditions are evaluated. The strongest stimulation is achieved when the target measure in the complete ROI exceeds the given threshold value (termed *t*_ROI_in the following). This corresponds to the maximally achievable *sensitivity* of 1, i.e., it maximizes the number of true positives in the ROI. Achieving the best focality requires that all elements in the non-ROI are kept below another specified threshold *t*_nonROI_. This corresponds to the highest *specificity* of 1, i.e., to no false positives in the non-ROI. The ROC curve graphically represents the relation between 1-*specificity* (*x*-axis) and the *sensitivity* (*y*-axis) of the target measure for varying threshold choices. A fully optimal solution corresponds to position (0,1) in the plot. By using the ROC, we aim to ensure a desired intensity-focality tradeoff and not just to maximize focality neglecting the intensity. The desired intensity-focality tradeoff can be chosen by setting the two thresholds for the target measure in the ROI and non-ROI. For TES, TIS and ECT, the goal function can then be defined by means of the Euclidean distance between the optimal position (0,1) and the point in the ROC plot that given by the achieved sensitivity and specificity of a solution.

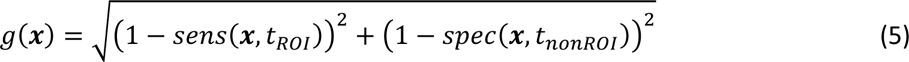

**Fig. 3:**
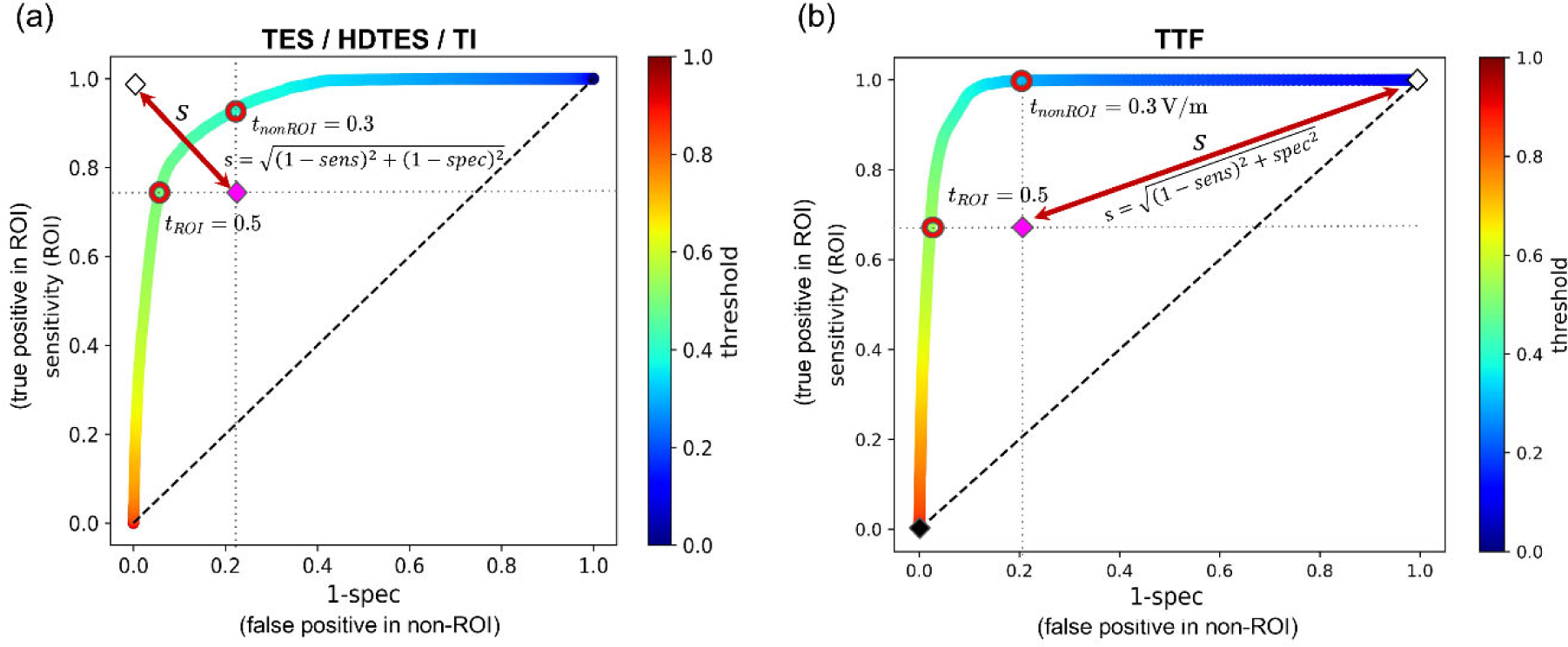
(a) Focality optimization: Optimization of the ROC curve; The focality is improved by *minimizing* the distance *s* between the reference location at (1 - *spec*, *sens*) = (0,1) and the current iteration. The optimization criterion was chosen to be particularly strict here by defining two separate thresholds such as the field should be greater than 0.5 (a.u.) in the ROI and smaller than 0.3 (a.u.) in the non-ROI; (b) Anti-focality optimization: Optimization of the ROC curve to improve the field *spread* for TTFields while ensuring high electric field values in the tumor region (ROI). Here it is the goal to *minimize* the distance *s* at (1 - *spec*, *sens*) = (1,1), which maximizes the electric field in the ROI and simultaneously makes the stimulation as unspecific as possible (i.e. make the number of false positives in the non-ROI as high as possible).

Interestingly, for TTFields, a clinically more relevant aim is to maximize the intensity in the ROI, while also maintaining high intensities in the rest of the brain (i.e. *minimizing* focality or *maximizing* “anti-focality”) in order to target both active tumor and diffusely infiltrating cancer cells (Korshoej et al., 2019b; Korshoej et al., 2020; Mikic et al., 2021; Mikic et al., 2024; Ballo et al., 2019). This corresponds to position (1,1) as being optimal in the ROC plot, and the corresponding distance-related goal function is then given as:

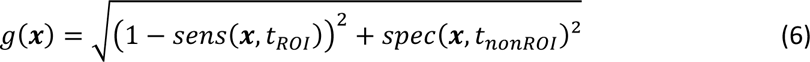

### 2.4. Optimization Approach

The optimization aims to determine the parameter vector 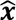, which comprises the center positions and orientations of the rectangular electrodes in case of standard TES, or the distances between the center and surround electrodes for 4×1 TES, that minimize the goal function *g*(**x**):

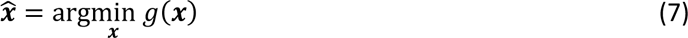

The parameters are subject to a number of constraints. Here, we ensure practically feasible solutions by allowing electrode positions only within a suited region of the upper head surface (Fig. 2b). Electrode configurations outside this area, partially overlapping with the border, or overlapping each other are penalized during the optimization. Setting further constraints for specific parameters is possible, such as defining the range of allowed distances between the center and surround electrodes for 4×1 TES. In combination, the variety of implemented goal functions, an intuitive description of the optimization problem by means of practically meaningful parameters, and relevant constraints provides a high degree of flexibility.

The optimization algorithm was chosen to give robust results also for TTFields, where many of the parameter combinations cause mutual overlaps of the large electrode arrays or arrays that fall partly outside of the upper skin region. This non-continuous solution space with numerous local minima, together with the properties of most of the goal functions defined above, makes the optimization non-convex and computationally demanding, even though the relatively low number of parameters (compared to lead-field based approaches) benefits the stability of the solution. For this reason, a stochastic optimization approach based on the differential evolution algorithm (Storn and Price, 1997) was chosen. Here, we used the `differential_evolution` algorithm implemented in *SciPy* (Virtanen et al., 2020), with a relative tolerance interval of 0.1 as convergence criterion. After optimization, a standard SimNIBS simulation with full electrode models is run to determine the final electric field distribution for the optimized parameters. Details on the hyperparameter choices are given in Supplementary Material S3, which also summarizes all steps of the overall approach.

### 2.5. Application Examples

The optimization algorithm was evaluated for standard two-electrode TES, focal 4×1 TES, TIS, and TTFields, using the *ernie* headmodel from the SimNIBS example dataset (https://simnibs.github.io/simnibs/build/html/dataset.html). The parameters for the field simulations and application examples are explained in more detail in the following.

#### 2.5.1. Head model

The *ernie* headmodel was created using the CHARM pipeline (Puonti et al., 2020) from SimNIBS v4.0 (Saturnino et al., 2019a) and consists of 9 different tissues given in Table 1. For TTFields, the tissue segmentation was manually modified to include an artificial subcortical residual located in the region of the right temporal lobe, together with a resection cavity. We defined application-specific regions of interests (ROI), where the electric field distribution is optimized. The ROIs are described in the following sub-sections together with the corresponding electrode setups.

**Table 1:**
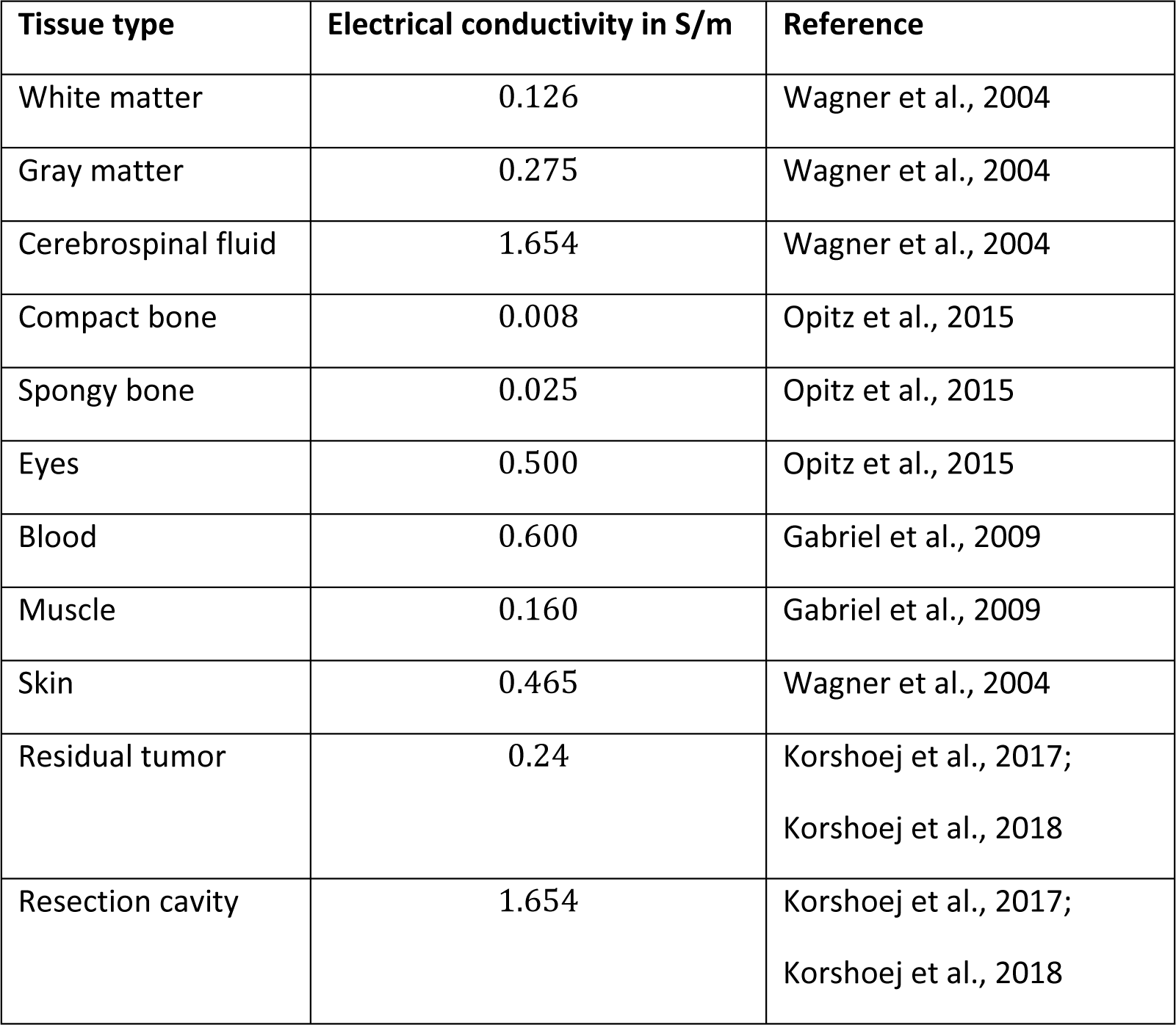
Tissue types and associated electrical conductivities of the headmodel.

#### 2.5.2. Standard Transcranial Electric Stimulation (TES)

The optimization algorithm was applied to a classic TES montage consisting of two large electrodes of rectangular shape (50 mm x 70 mm; Fig. 4a). The first goal was to determine the electrode configuration that generates the highest average electric field strength in the motor cortex (M1), using eq. (3) as goal function. In a second optimization run, the aim was to optimize the focality of the electric field in the motor cortex according to eq. (5) as goal function (Fig. 3a). Two position parameters and one orientation parameter per electrode were optimized, resulting in a total of six free parameters. A current of *I*_max_ = 2 mA was assumed for both cases and the thresholds for focality optimization were *t*_ROI_ = 0.2 V⁄m and *t*_nonROI_ = 0.1 V⁄m. The gray matter midlayer surface of the handknob region of M1 was defined as the ROI (green in Fig. 4a). For focality optimization, the remaining surface of the midlayer was defined as the non-ROI.

**Fig. 4:**
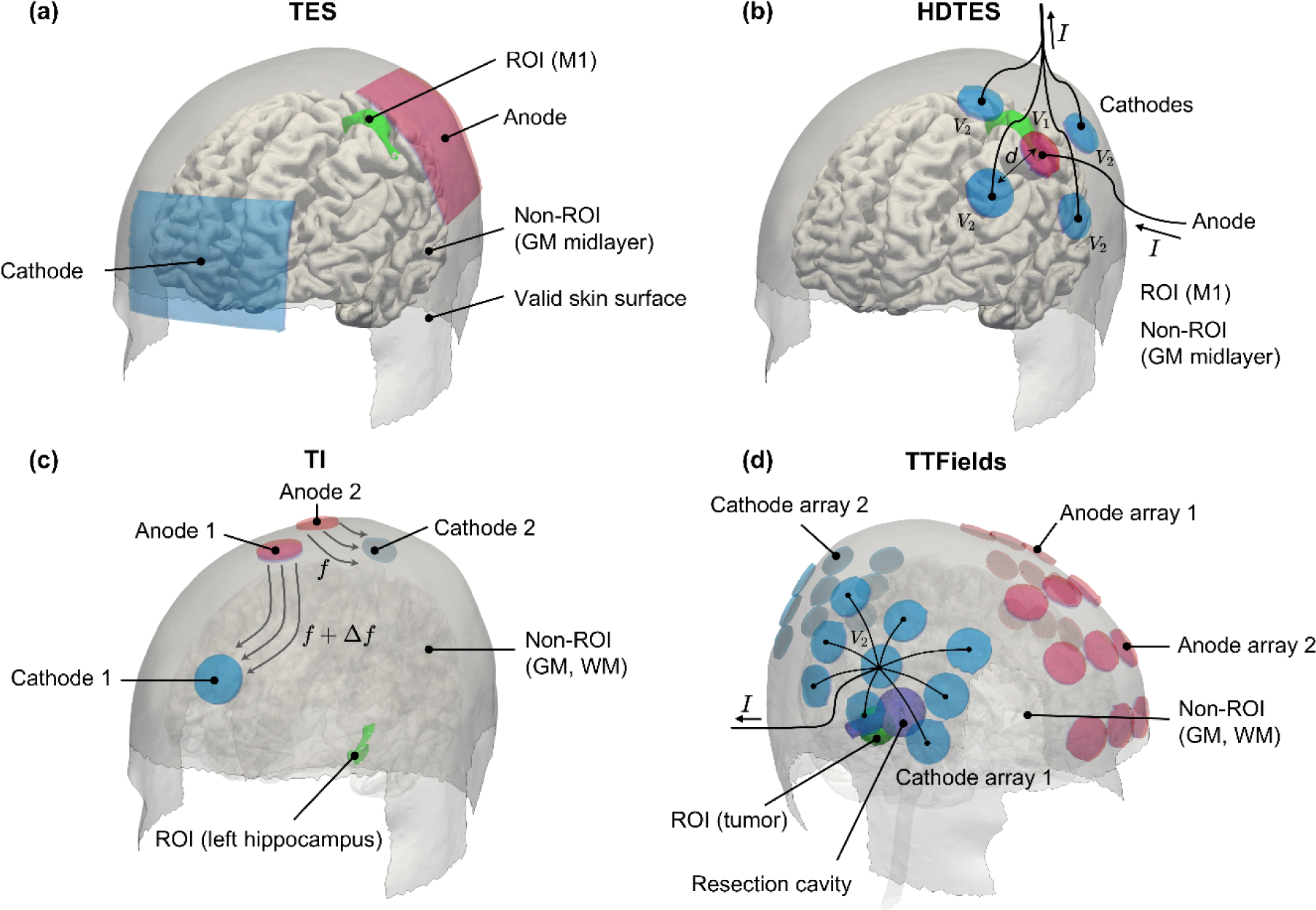
Application examples: (a) Standard Transcranial Electric Stimulation (TES) using two large electrode patches; (b) Focal 4×1 TES using one inner electrode and four outer electrodes located at variable distance *d*; (c) Temporal interference stimulation (TIS) using two pairs of small electrodes impressing currents with different frequencies; (d) Tumor Treating Fields (TTFields) using two pairs of 3×3 electrode arrays.

The optimization was performed using the *node-wise Dirichlet correction* to account for the non-uniform current density distribution at the electrode-skin interface. The influence of neglecting this correction was also investigated. In this case, a constant current density was impressed over the entire electrode surface. This simplification accelerates the optimization. It was investigated to what extent this choice affects the final value of the goal function and the computing time, in order to assess whether applying the correction is in fact necessary and worthwhile.

#### 2.5.3. Focal multi-electrode Transcranial Electric Stimulation (Focal 4×1 TES)

The second example targeted the optimization of a focal 4×1 TES montage (Fig. 4b), consisting of circular electrodes with diameters of 20 mm. The currents and thresholds were the same as in the standard TES case described above. The inner electrode was connected to the first channel, and the four outer electrodes commonly to the second channel. The optimization variables were two parameters for the position of the inner electrode, and one orientation and one distance parameter to describe the positions of the outer electrodes relative to the inner one, resulting in four free variables. The distance between inner and outer electrodes was restricted to the interval [25, 100] mm. Here again, the aim was to find two electrode configurations, where the first maximizes the mean electric field strength in the motor cortex (ROI), while the second optimizes the focality.

The effect created by several individual electrodes connected to one channel can be accounted for by applying the *electrode-wise Dirichlet correction*, which can be interpreted as an intermediate solution between no correction and the exact *node-wise Dirichlet correction*. It ensures the correct total currents for each of the four outer electrodes (see *I*_l_… *I*_4_ in Fig. 1), which is computationally far less demanding than correcting the currents in each skin node related to one of the outer electrodes. The influence of the correction type (*electrode-wise, node-wise, no correction)* on the electric field, the optimization result, and the computing time were investigated to determine the best setting for future use.

#### 2.5.4. Temporal interference stimulation (TIS)

The third example addressed the optimization of a TIS montage (Fig. 4c). The setup consisted of two pairs of circular electrodes with a diameter of 20 mm each. The currents and thresholds were the same as in the TES cases described above. The goal was to optimize the electric field envelope according to eq. (1) in the left hippocampus defined as the ROI, either in terms of intensity or focality. The remaining grey and white matter volumes were defined as the non-ROI. For TIS, evaluation of the goal function required two electric field calculations per iteration, one for each channel, which were superimposed to calculate the maximum amplitude of the envelope according to eq. (1). Afterwards, the goal function value, depending on the optimization problem, was calculated. Given four electrodes with two position parameters each, the total number of free variables was eight. The influence of applying *node-wise Dirichlet correction* on the solution was tested, as applying no correction and assuming a constant current density in the small electrodes accelerates the optimization.

#### 2.5.5. Tumor Treating Fields (TTFields)

The final example targeted the optimization of a TTFields montage (Segar et al., 2023) that consists of two pairs of electrode arrays (Fig. 4d). Each array comprises a three-by-three layout of circular electrodes that are spaced vertically by 22 mm and horizontally by 33 mm, and have diameters of 20 mm. The *ernie* head model was modified by adding tumor tissue, necrosis, and surrounding edema. The tumor volume was defined as the ROI and the remaining gray and white matter was defined as the non-ROI. The first optimization goal was to find an electrode arrangement, which maximizes the field intensity in the tumor according to eq. (3) as goal function. In a second optimization, it was the goal to create strong field intensities in the tumor, which are above a given threshold, while maximizing the field exposure also in the remaining brain. For that, the goal function of each array pair was calculated according to eq. (6) (Fig. 3b) and the resulting values of both pairs combined using eq. (4). This requires two electric field calculations, one per array pair, per iteration. The total current in each array pair was defined to *I*_max_ = 1 A baseline-to-peak and the thresholds were *t*_ROI_ = *t*_nonROI_ = 150 V⁄m. Each of the four electrode arrays was described by two position parameters and one orientation parameter, resulting in a total of 12 free variables.

The choice of the correction type (*electrode-wise, node-wise, no correction*) on the speed and accuracy of the solution was also tested. As stated further above, the non-continuous solution space and the moderately higher number of free variables compared to the other examples makes TTFields the most demanding optimization problem tested here. Mathematically, this is represented by an enlargement and clustering of the hyperdimensional constraint surface, which is superimposing the goal function, making it challenging to find an optimal solution. This generally resulted in a higher number of function evaluations, and also caused a strong impact of the chosen correction type on the duration of the optimization.

#### 2.5.6. Performance evaluation

To enable an evaluation of the performance of the optimization procedure and its dependence on the chosen current correction type, 200 valid parameter sets were randomly created for each of the above examples. For each set, the results for the three correction types and additionally for a standard SimNIBS simulation with full electrode models were obtained. To assess the impact of the correction type on the simulated fields, the fields for the three correction types in gray and white matter were correlated with the field of the standard simulation, and a mean correlation coefficient and the standard deviation were determined over the 200 electrode positions. Normalized root-mean-square deviations (NRMSD) between the fields were additionally calculated and are reported in the Supplemental Material Section S5.

In addition, the respective goal function values for intensity and focality were determined from the standard SimNIBS results of the random parameter sets to get reference distributions for assessment of the optimized solution. As the selected optimization algorithm *differential evolution* is stochastic, the optimization results may vary between two repetitions. Therefore, 30 independent repetitions of the optimization were carried out for each application, and the resulting goal function values compared to the reference distributions from the random parameter sets. The optimizations were repeated for the different current correction approaches to investigate their influence on the result.

The simulations were performed on a Ryzen 9 5950X CPU with 3.4 GHz (16 cores, 128 GB RAM) and the computing time is recorded for comparison.

## 3. Results

### 3.1. Standard Transcranial Electric Stimulation (TES)

In order to evaluate the effect of the chosen current correction type, the correlation coefficients of the electric fields and goal function values obtained for node-wise current corrections and no corrections, respectively, with the values for the reference simulations are shown in Table 2. As expected, the mean correlation coefficients are lower when no corrections are applied. The results also show that this affects the focality goal function more than the intensity goal function. The distributions of the NRMSD over the 200 simulations are shown in Supplementary Fig. S3a and confirm the differences in accuracy between the two correction approaches. Differences in the simulated electric fields will cause differences in the goal functions values, which can impact the quality of the optimization results. However, it is worth noting that the absolute values of the electric fields play a secondary role for the optimization and it is more important that relative changes across different parameter sets are mapped correctly. As long as relative changes are represented accurately, a more efficient modeling approach will lead to the same final parameter set and can thus be used during optimization. The final results will then not differ, as a standard SimNIBS simulation is carried out as last step in any case.

**Table 2:**
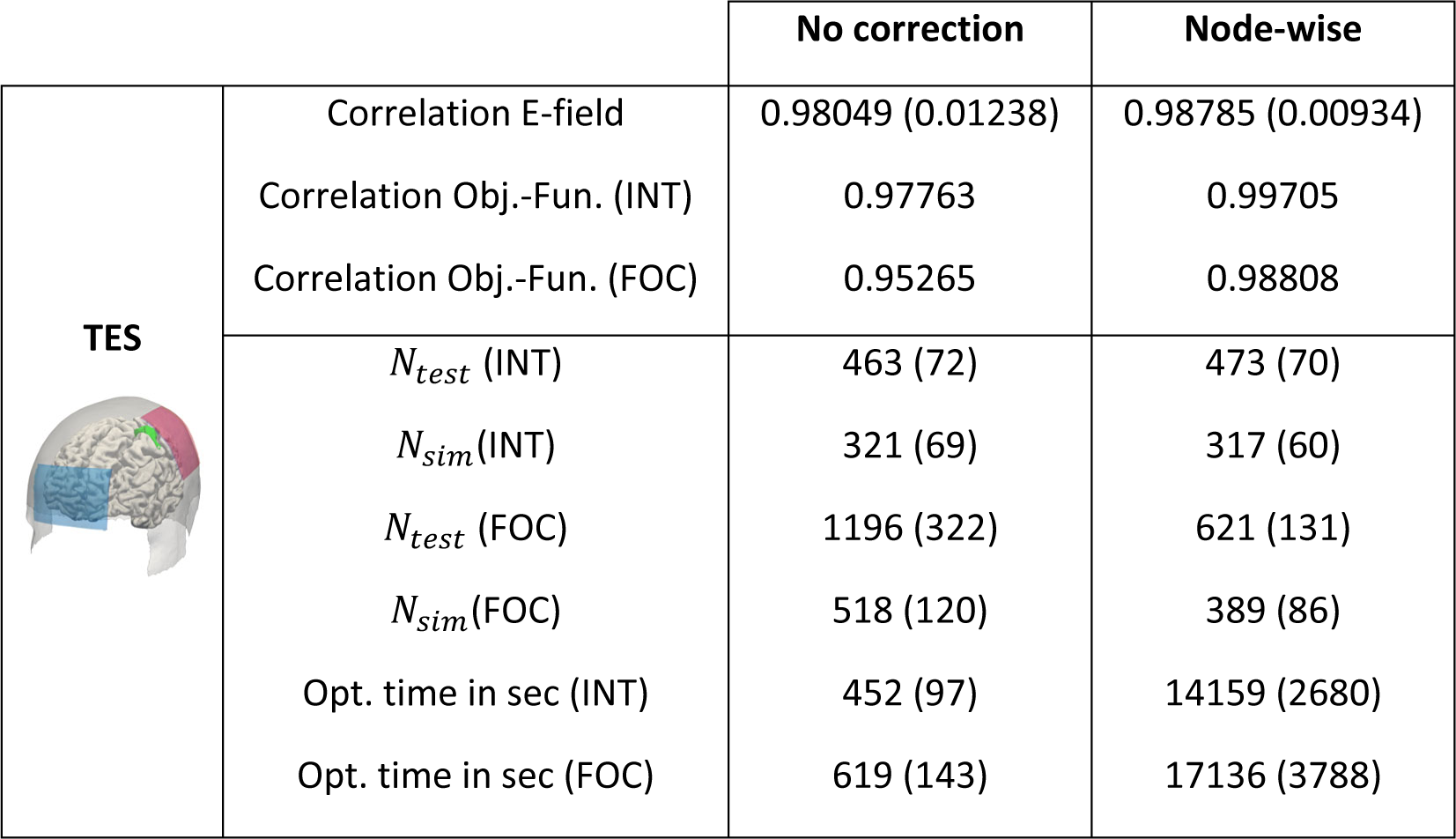
Summary results for conventional TES. Upper part: Comparison between the results for the different current correction methods to the reference simulations over 200 repetitions. Average correlation coefficients of the electric field and the respective goal functions values for intensity (INT) and focality (FOC) optimization are provided. Lower part: Assessment of the efficiency of optimization procedure. *N*_test_ and *N*_sim_ denote the number of electrode placement tests and actual electric field simulations, respectively, averaged over 30 repeated optimization runs. The total computation time of the optimization runs are given in the last two rows. The standard deviations are given in brackets.

|  |  | No correction | Node-wise |
| --- | --- | --- | --- |
| **TES** | Correlation E-field | 0.98049 (0.01238) | 0.98785 (0.00934) |
|  | Correlation Obj.-Fun. (INT) | 0.97763 | 0.99705 |
|  | Correlation Obj.-Fun. (FOC) | 0.95265 | 0.98808 |
| | $N_{test}$ (INT) | 463 (72) | 473 (70) |
| | $N_{sim}$(INT) | 321 (69) | 317 (60) |
| | $N_{test}$ (FOC) | 1196 (322) | 621 (131) |
| | $N_{sim}$(FOC) | 518 (120) | 389 (86) |
|  | Opt. time in sec (INT) | 452 (97) | 14159 (2680) |
|  | Opt. time in sec (FOC) | 619 (143) | 17136 (3788) |

Fig. 5a shows a representative example of an optimized electric field, aimed at maximizing the electric field intensity in the M1 ROI. The effects of the node-wise current correction are clearly visible as inhomogeneous current distribution at the electrode edges. The histograms of the objective function values of the 30 repetitions of the optimization and the 200 random parameter sets are shown in Fig. 5b. The optimizations both with node-wise and no current corrections achieved consistently high goal function values, suggesting that using no current corrections to speed up the optimization is feasible for this application.

**Fig. 5:**
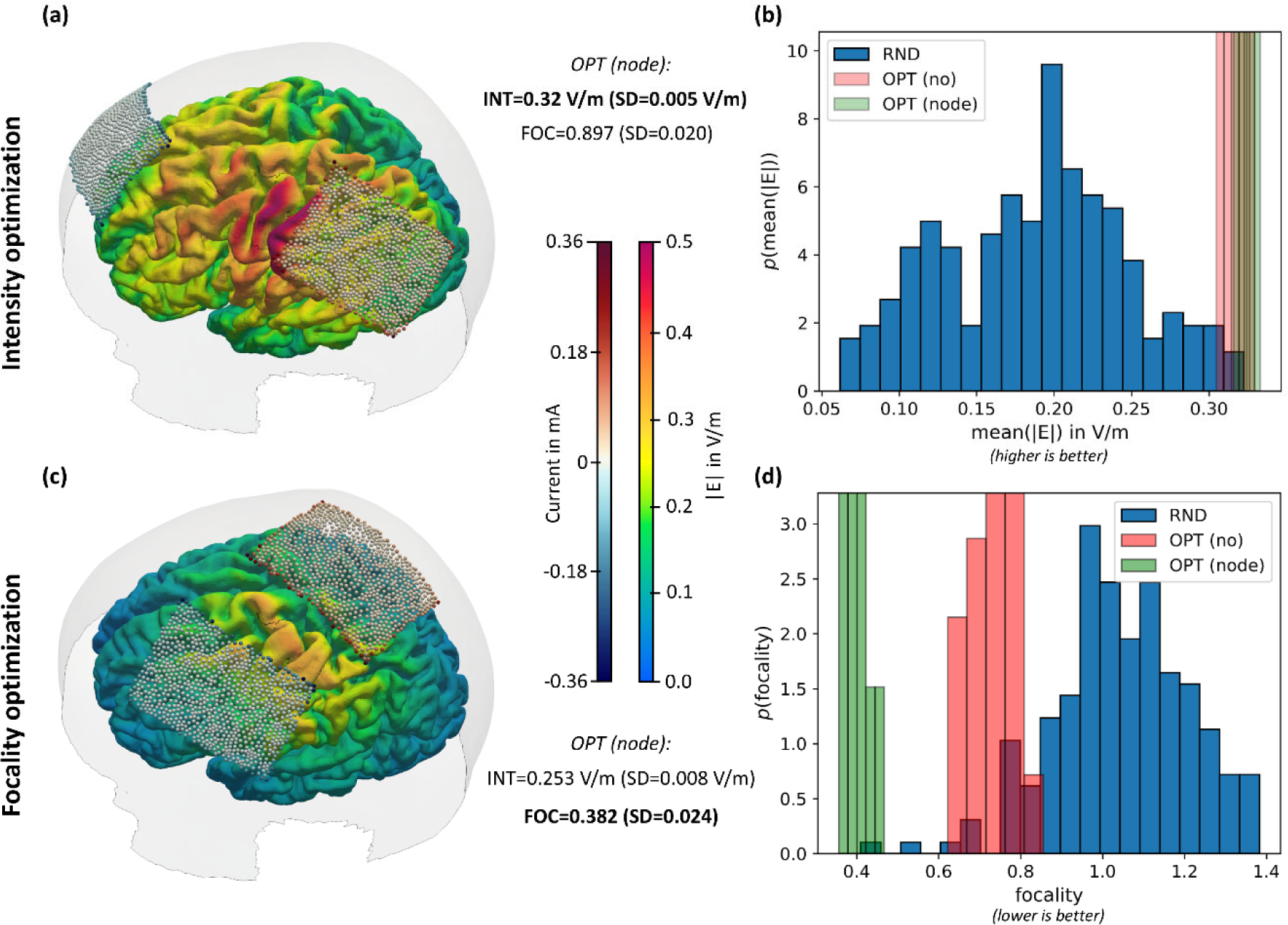
Optimization results for conventional TES. (a) Representative example of intensity based optimization showing the resulting electric field in the brain; (b) Histograms of the average electric field magnitude (higher is better) determined from 200 random electrode configurations (RND) and 30 optimization runs (OPT). The optimizations were performed without Dirichlet correction (no) and with node-wise Dirichlet correction (node), but the shown objective function values were determined in final reference simulations; (c) Representative example of focality based optimization; (d) same as (b) but goal function is focality according to distance *s* in Fig. 3(a) (lower is better).

In contrast, when optimizing the focality (Fig. 5c&d), using no current corrections results in worse final goal function values compared to node-wise corrections, despite of all optimizations completing successfully. This suggests that the (inhomogeneous) field distribution at the skin interface changes too much with the tested parameters, also affecting the goal function value strongly enough to impact the optimization result. Thus, applying node-wise current corrections is necessary in this case.

Comparing both optimization results illustrates the differences between intensity- and focality-based goal functions clearly. A substantially higher electric field can be generated by placing the electrodes further apart, at the expense of focality. On the other hand, placing the electrodes closer together increases focality, while decreasing the field strength in the ROI.

During optimization, it occurs that the selected parameters cause the electrodes to mutually overlap or being (partly) outside the permitted skin mask. The process is truncated in those cases, no electrical field calculation is performed, and the objective function is penalized. This reduces the calculation time in this iteration. The total number of times where an attempt was made to place the electrodes is labeled *N*_test_ in Table 2. The number of successful placements that are followed by field calculations is labeled is labeled *N*_sim_.

### 3.2. Focal 4×1 TES

The optimization results for 4×1 TES are summarized in Fig. 6 and Table 3. As the electrodes are considerably smaller than for standard TES, inhomogeneous field distributions at electrode-skin interfaces do hardly affect the electric field in the brain. The electrode-wise current correction that accounts for the effect caused by the outer electrodes being connected to the same channel thus achieves fields that are highly similar to those of a full simulation (Table 3). Interestingly, this also holds when using no correction, in which case it is assumed that the return currents through the outer electrodes are the same, i.e. *I*_l_ = ⋯ = *I*_4_ = *I*⁄4. This suggests that the volume conductivities between the inner electrode and each of the outer electrodes are comparable.

**Fig. 6:**
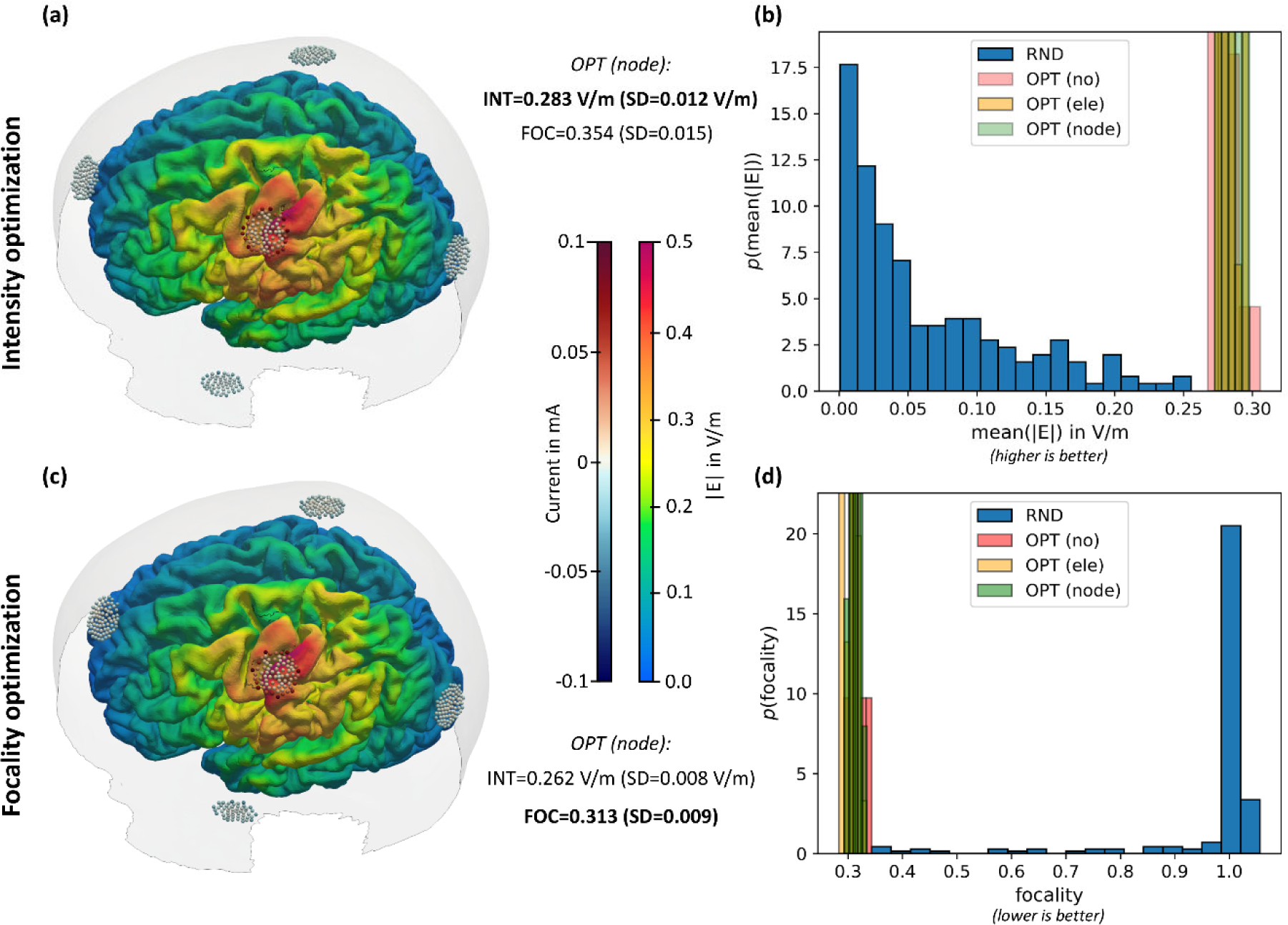
Optimization results for focal 4×1 TES. (a) Representative example of intensity based optimization showing the resulting electric field in the brain; (b) Histograms of the average electric field magnitude (higher is better) determined from 200 random electrode configurations (RND) and 30 optimization runs (OPT). The optimizations were performed without Dirichlet correction (no) and with node-wise Dirichlet correction (node), but the shown objective function values were determined in final reference simulations; (c) Representative example of focality based optimization; (d) same as (b) but goal function is focality according to distance *s* in Fig. 3(a) (lower is better).

**Table 3:**
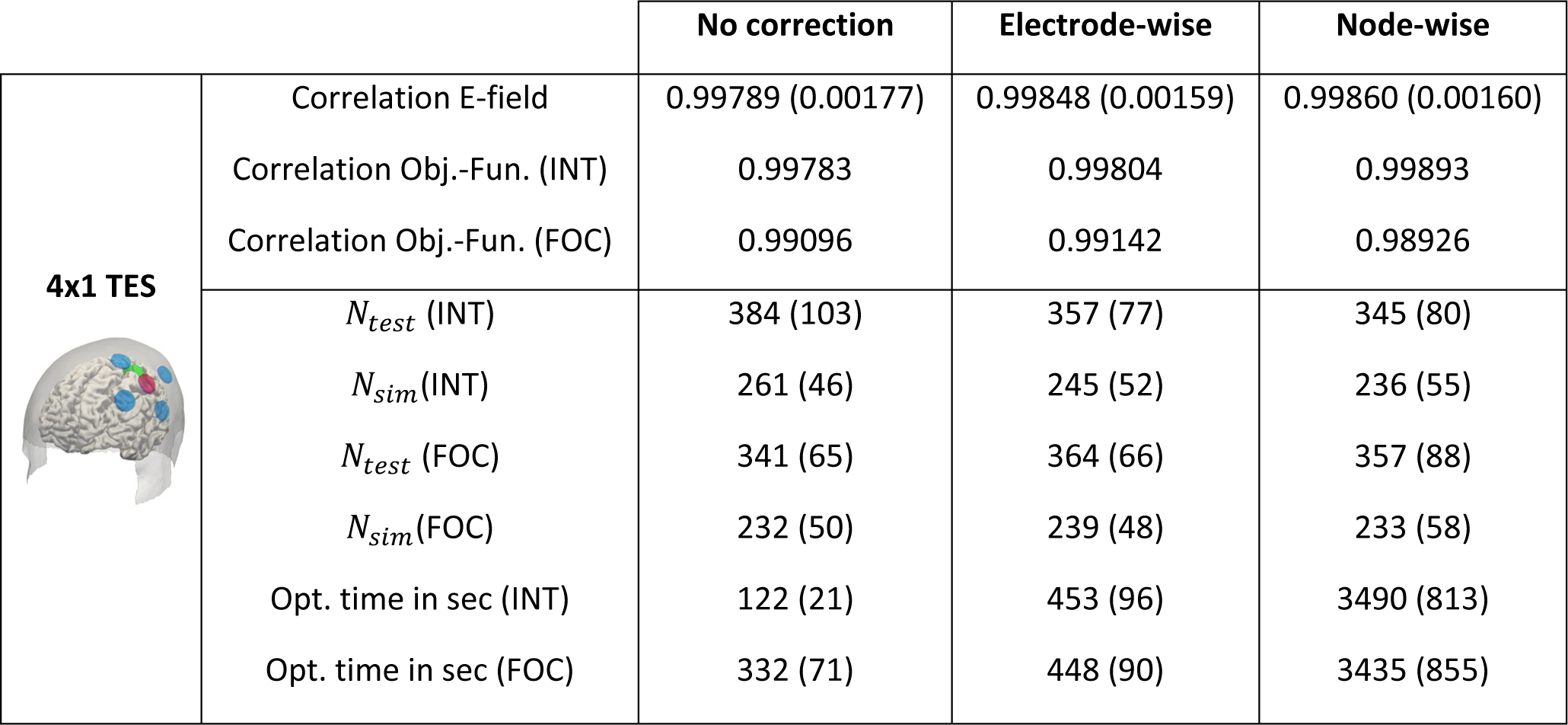
Summary results for focal 4×1 TES. Upper part: Comparison between the results for the different current correction methods to the reference simulations over 200 repetitions. Average correlation coefficients of the electric field and the respective goal functions values for intensity (INT) and focality (FOC) optimization are provided. Lower part: Assessment of the efficiency of optimization procedure. *N*_test_ and *N*_sim_ denote the number of electrode placement tests and actual electric field simulations, respectively, averaged over 30 repeated optimization runs. The total computation time of the optimization runs are given in the last two rows. The standard deviations are given in brackets.

|  |  | No correction | Electrode-wise | Node-wise |
| --- | --- | --- | --- | --- |
| **4x1 TES** | Correlation E-field | 0.99789 (0.00177) | 0.99848 (0.00159) | 0.99860 (0.00160) |
|  | Correlation Obj.-Fun. (INT) | 0.99783 | 0.99804 | 0.99893 |
|  | Correlation Obj.-Fun. (FOC) | 0.99096 | 0.99142 | 0.98926 |
| | $N_{test}$ (INT) | 384 (103) | 357 (77) | 345 (80) |
| | $N_{sim}$(INT) | 261 (46) | 245 (52) | 236 (55) |
| | $N_{test}$ (FOC) | 341 (65) | 364 (66) | 357 (88) |
| | $N_{sim}$(FOC) | 232 (50) | 239 (48) | 233 (58) |
|  | Opt. time in sec (INT) | 122 (21) | 453 (96) | 3490 (813) |
|  | Opt. time in sec (FOC) | 332 (71) | 448 (90) | 3435 (855) |

As result, the optimization results shown in Fig. 6 robustly yield high goal function values, irrespective of the chosen current correction approach. Generally, we suggest applying no corrections. Alternatively, to ensure reliable results also in cases where the volume conduction properties vary more strongly across electrode positions, e.g. in patients with cranial openings, electrode-wise corrections will represent an efficient compromise between accuracy and calculation time.

### 3.3. Temporal Interference Stimulation (TIS)

The results for TIS are shown in Figure 7 and Table 4. Given the small electrode sizes, it is expected that the chosen current correction method (no corrections, node-wise corrections) does not affect the optimization results, so that applying no corrections is feasible to speed up the optimization.

**Fig. 7:**
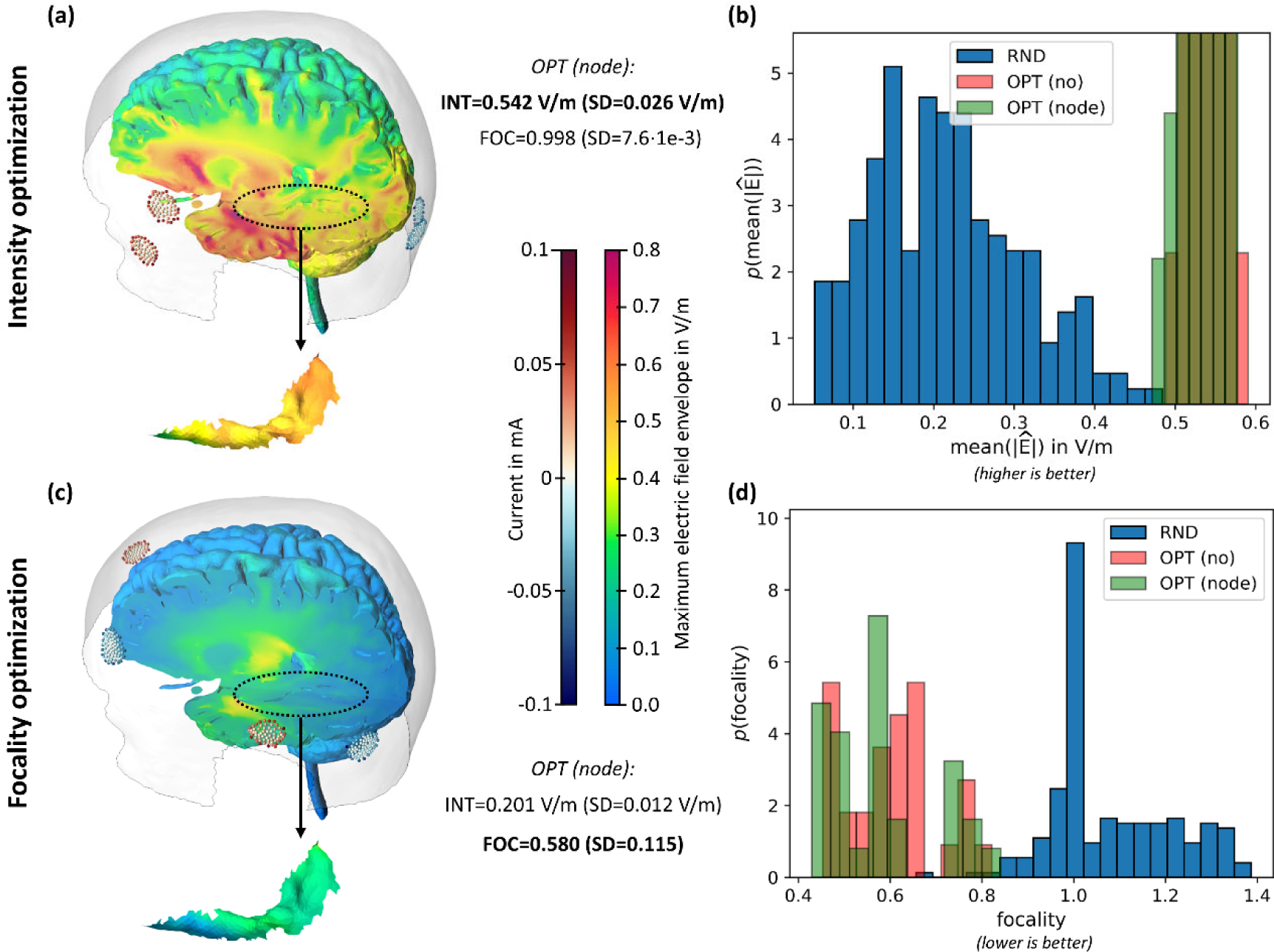
Optimization results for Temporal Interference Stimulation (TIS). (a) Representative example of intensity based optimization showing the resulting maximum electric field envelope in the brain; (b) Histograms of the average maximum electric field envelope (higher is better) determined from 200 random electrode configurations (RND) and 30 optimization runs (OPT). The optimizations were performed without Dirichlet correction (no) and with node-wise Dirichlet correction (node), but the shown objective function values were determined in final reference simulations; (c) Representative example of focality based optimization; (d) same as (b) but goal function is focality according to distance *s* in Fig. 3(a) (lower is better).

**Table 4:**
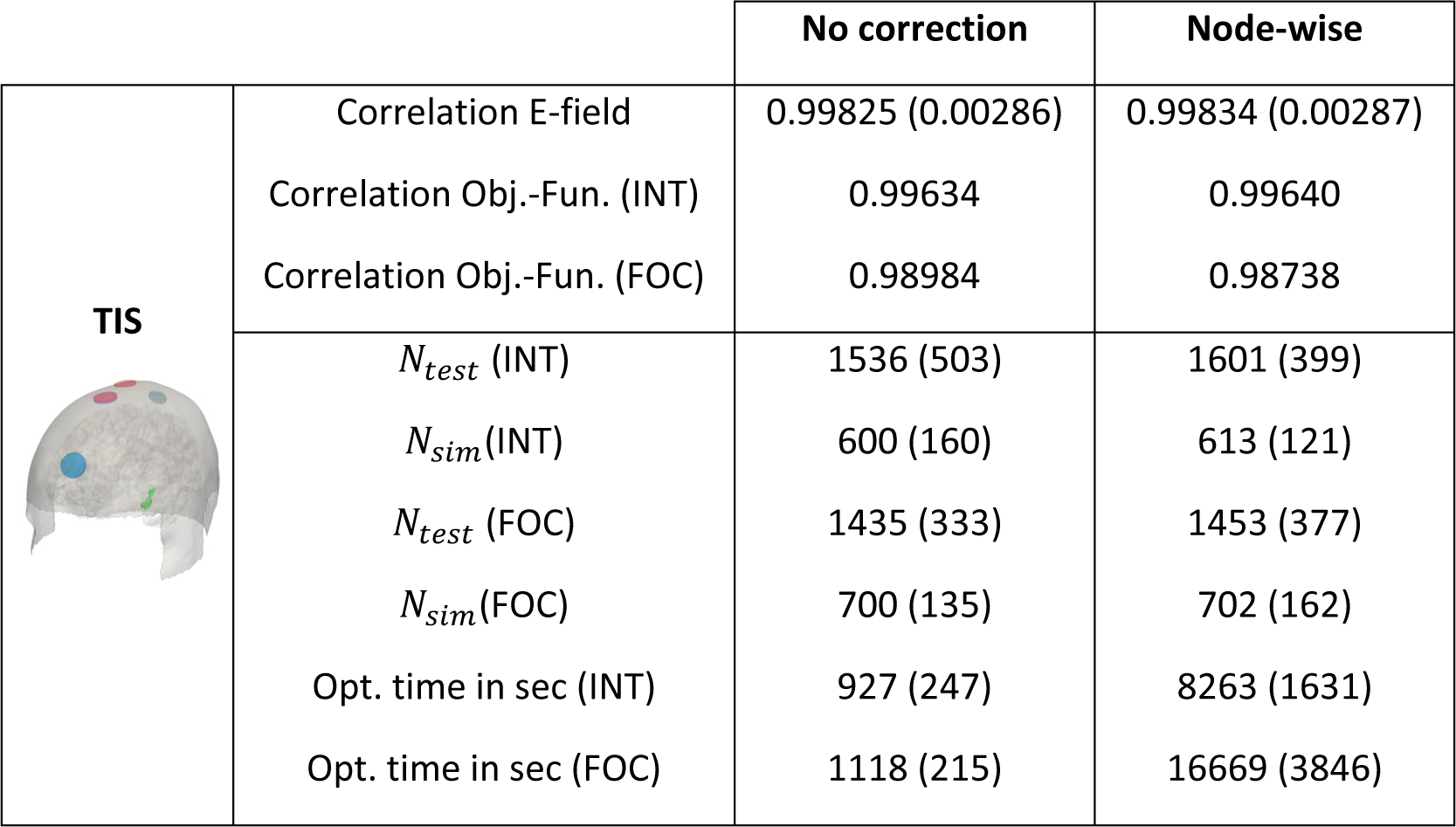
Summary results for TIS. Upper part: Comparison between the results for the different current correction methods to the reference simulations over 200 repetitions. Average correlation coefficients of the electric field and the respective goal functions values for intensity (INT) and focality (FOC) optimization are provided. Lower part: Assessment of the efficiency of optimization procedure. *N*_test_and *N*_sim_ denote the number of electrode placement tests and actual electric field simulations, respectively, averaged over 30 repeated optimization runs. The total computation time of the optimization runs are given in the last two rows. The standard deviations are given in brackets.

|  |  | No correction | Node-wise |
| --- | --- | --- | --- |
| **TIS** | Correlation E-field | 0.99825 (0.00286) | 0.99834 (0.00287) |
|  | Correlation Obj.-Fun. (INT) | 0.99634 | 0.99640 |
|  | Correlation Obj.-Fun. (FOC) | 0.98984 | 0.98738 |
| | $N_{test}$ (INT) | 1536 (503) | 1601 (399) |
| | $N_{sim}$(INT) | 600 (160) | 613 (121) |
| | $N_{test}$ (FOC) | 1435 (333) | 1453 (377) |
| | $N_{sim}$(FOC) | 700 (135) | 702 (162) |
|  | Opt. time in sec (INT) | 927 (247) | 8263 (1631) |
|  | Opt. time in sec (FOC) | 1118 (215) | 16669 (3846) |

Interestingly, the achieved goal function values when optimizing the focality for TIS (Fig. 7c&d) show a larger spread across the 30 repeated optimization runs, compared to the other tested applications. This suggests that focality optimization for TIS is more challenging, likely due to the more complex goal function that is based on the non-linear combination of two superimposed electric fields, which are then fed into the ROC-based evaluation of the focality-intensity tradeoff (combining eq. 1 & 5). This increases the probability that the optimization algorithm converges to a local minimum. In practice, a multi-start approach can, however, be used to prevent this.

### 3.4. Tumor Treating Fields (TTFields)

The results for TTFields are shown in Figure 8 and Table 5. As all electrodes of an array share the same channel, also the electrode-wise current correction is tested again. All current correction approaches result in electric fields and goal function values that are very similar to those of the reference simulations (Table 5). In consequence, also their final optimized parameter sets are similar (Fig. 8), so that running the optimization without current correction is feasible to minimize the required time. Alternatively, using electrode-wise current correction will help to ensure reliable results also, e.g., in patients with cranial openings.

**Fig. 8:**
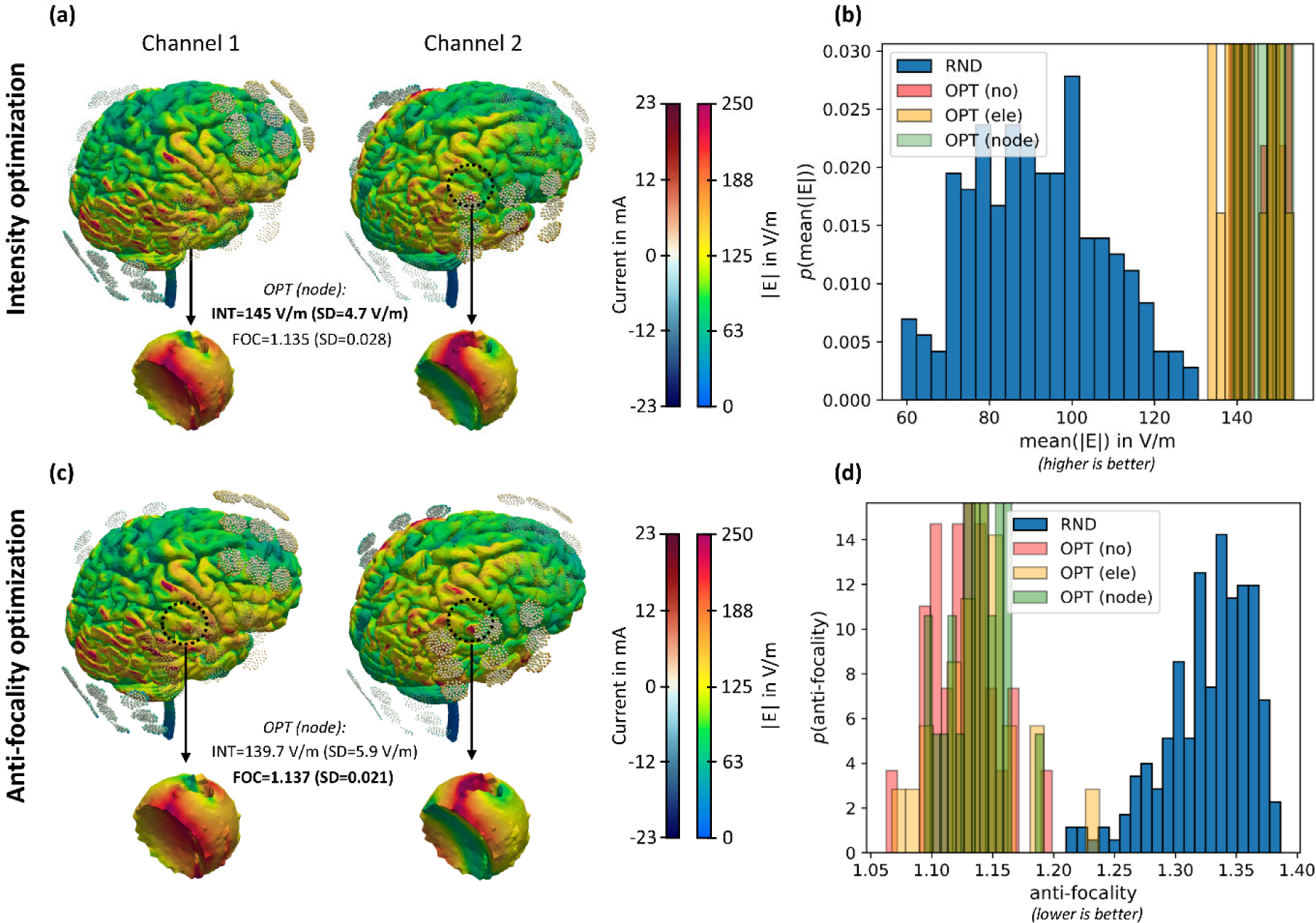
Optimization results of Tumor Treating Fields (TTFields). (a) Representative example of intensity-based optimization showing the resulting electric field in the brain; (b) Histograms of the average electric field magnitude (higher is better) determined from 200 random electrode configurations (RND) and 30 optimization runs (OPT). The optimizations were performed without current correction (no) and with node-wise current correction (node), but the shown goal function values were determined in final reference simulations; (c) Representative example of anti-focality based optimization; (d) same as (b) but the goal function is anti-focality according to distance *s* in Fig. 3(b) (lower is better).

**Table 5:**
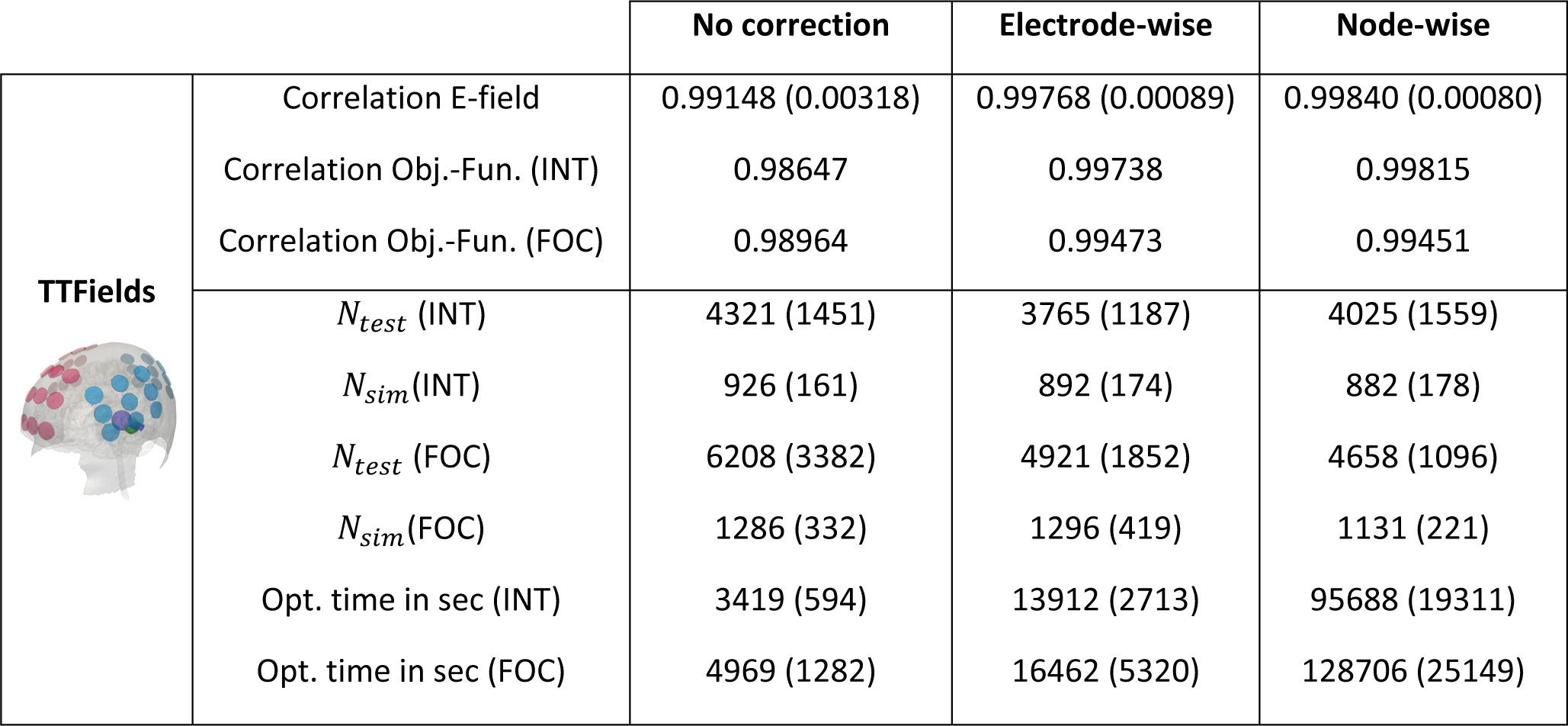
Summary results for TTFields. Upper part: Comparison between the results for the different current correction methods to the reference simulations over 200 repetitions. Average correlation coefficients of the electric field and the respective goal functions values for intensity (INT) and focality (FOC) optimization are provided. Lower part: Assessment of the efficiency of optimization procedure. *N*_test_and *N*_sim_ denote the number of electrode placement tests and actual electric field simulations, respectively, averaged over 30 repeated optimization runs. The total computation time of the optimization runs are given in the last two rows. The standard deviations are given in brackets.

|  |  | No correction | Electrode-wise | Node-wise |
| --- | --- | --- | --- | --- |
| **TTFields** | Correlation E-field | 0.99148 (0.00318) | 0.99768 (0.00089) | 0.99840 (0.00080) |
|  | Correlation Obj.-Fun. (INT) | 0.98647 | 0.99738 | 0.99815 |
|  | Correlation Obj.-Fun. (FOC) | 0.98964 | 0.99473 | 0.99451 |
| | $N_{test}$ (INT) | 4321 (1451) | 3765 (1187) | 4025 (1559) |
| | $N_{sim}$ (INT) | 926 (161) | 892 (174) | 882 (178) |
| | $N_{test}$ (FOC) | 6208 (3382) | 4921 (1852) | 4658 (1096) |
| | $N_{sim}$ (FOC) | 1286 (332) | 1296 (419) | 1131 (221) |
|  | Opt. time in sec (INT) | 3419 (594) | 13912 (2713) | 95688 (19311) |
|  | Opt. time in sec (FOC) | 4969 (1282) | 16462 (5320) | 128706 (25149) |

Due to the particularly large electrode arrays, this optimization problem is highly constrained. This increases the number of tested parameter sets *N*_test_ in Table 5 compared to the other stimulation methods. As a result, the relative number of effectively performed electric field simulations is also lower. Despite the challenging constraints, both intensity- and anti-focality-based optimizations consistently reach high goal function values for the 30 repetitions. However, due to the clinically sensitive application in tumor patients, we recommend a multi-start approach for TTFields to ensure best possible optimization results.

### 3.5. Further showcases for future applications

#### Geometry optimization of TES electrodes

In order to demonstrate the potential of the developed optimization approach, we optimized also the sizes of the two electrodes of a standard TES montage in addition to their positions and orientations. The currents and thresholds were the same as in the standard TES case described above. The edge length of the rectangular electrodes was allowed to vary between 10 and 70 mm. The results are shown in Fig. 9 for intensity and focality optimization. Interestingly, the optimization resulted in smaller electrodes being theoretically advantageous. Compared to the standard size, the mean electric field in the ROI could be increased by an additional 19% and the focality measure could also be improved by a considerable 56% compared to the optimal solutions shown in Fig. 5.

**Fig. 9:**
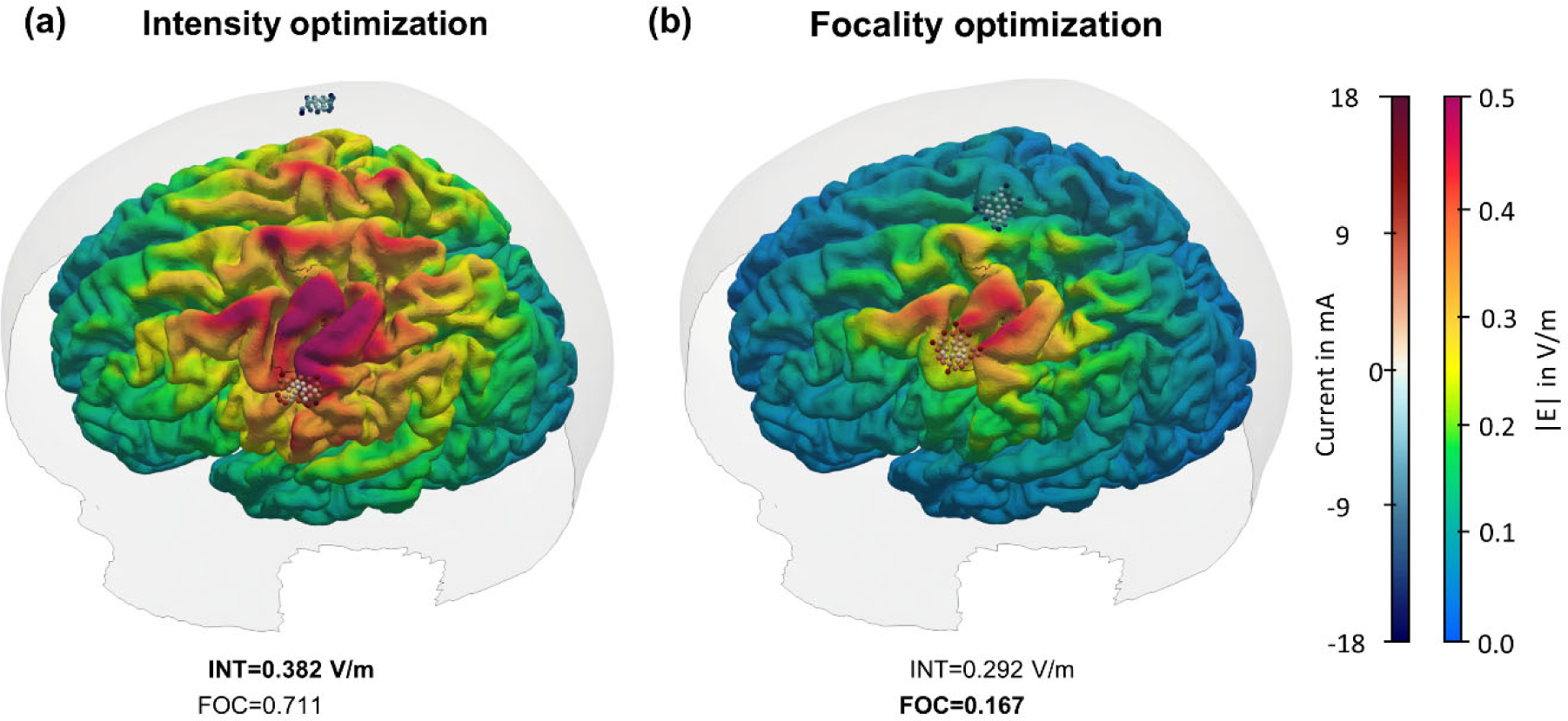
Optimization results of two-electrode TES including electrode size. (a) Intensity-based optimization and (b) focality based optimization showing the resulting electric field in the brain and the optimized sizes and locations of the rectangular electrodes, respectively.

#### Electroconvulsive therapy (ECT)

As a second use case, we optimized the electrode positions for ECT and compared it with the standard RUL montage. The radius of the round electrodes was 25 mm and the stimulation intensity was 800 mA. The right prefrontal cortex was defined as the ROI. To reduce adverse side effects, the left and right hippocampus were defined as non-ROI. The thresholds for the focality optimization were *t*_ROI_ = 120 V⁄m and *t*_nonROI_ = 50 V⁄m. The results of the intensity- and focality-optimal montages compared to standard RUL are shown in Fig. 10. The intensity of the average electric field in the right prefrontal cortex was increased in both cases from 86 V/m using the standard RUL montage to 113 V/m (+31%) for both intensity and focality optimization. The focality measure of the RUL montage was 0.51 and was increased to 0.82 for intensity optimization but was effectively reduced to 0.47 in case of focality optimization. It should be noted that the average electric field in the hippocampus increased from 52.2 V/m to 57.8 V/m (+11%) for intensity optimization and reduced to 44.5 V/m (−15%) for focality optimization respectively compared to the RUL montage.

**Fig. 10:**
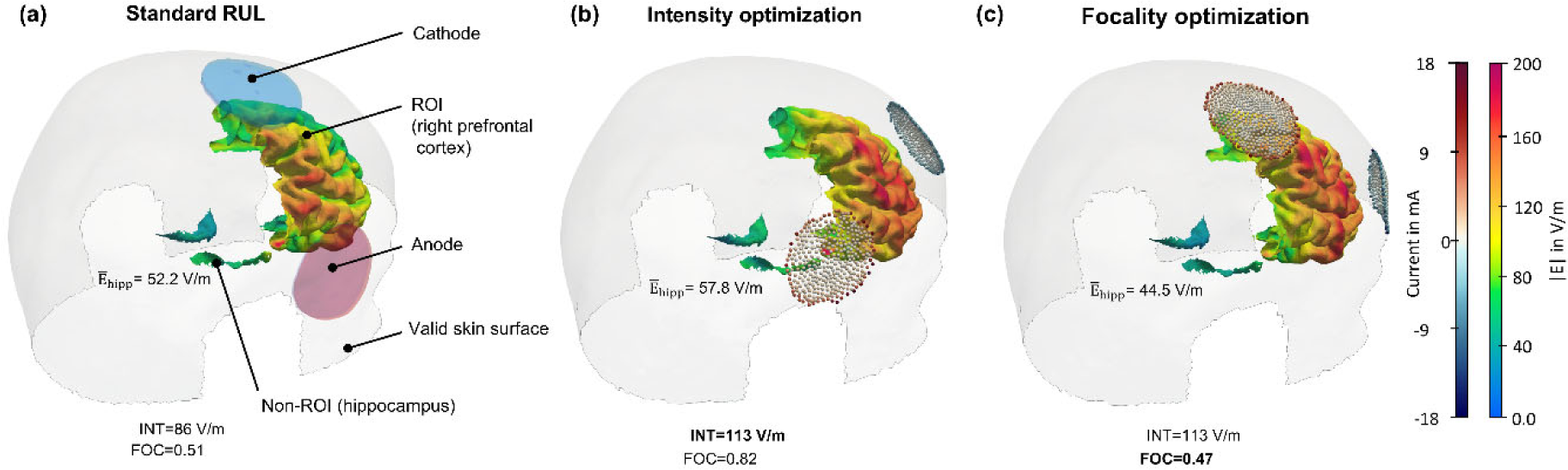
Optimization results of Electroconvulsive Therapy (ECT). (a) Standard RUL montage, (b) intensity-based optimization, and (c) focality based optimization showing the resulting electric field in the brain and the optimized electrode locations.

## 4. Discussion

### Summary of work

We developed and validated a flexible framework that enables the optimization of electrode configurations for different electric stimulation and treatment modalities, including standard TES, focal 4×1 TES, TIS, ECT and TTFields. The electrodes can move freely over the head surface independent of a predefined discretization and the approach accounts for spatial relationships between neighboring electrodes, as in 4×1 TES, as well as spatially extended electrode shapes, as in standard TES. As such, it serves an important complementary role to existing optimization approaches for multi-channel stimulation (Dmochowski et al., 2011; Fernandez-Corazza et al., 2020; Saturnino et al. 2019) that were designed for use with small electrodes placed at discrete positions and require multi-channel stimulation systems.

Our new optimization approach supports intensity-based optimizations to maximize the target measure in the ROI, as well as ROC-based optimizations for a flexible control of the tradeoff between intensity and focality, based on the threshold values chosen for the ROI and non-ROI. Various target measures such as the magnitude or normal component of the electric field and the TIS field envelope are supported. The provided examples cover multiple relevant clinical and scientific application scenarios with diverse constraints. In order to support the integration into practical workflows, we put emphasis on an efficient implementation that keeps the system requirements moderate and the computing times low enough for use on standard computers.

### Quality and robustness of the optimization results

The optimization results can be affected by the accuracy of the approximation of the inhomogeneous current flow distribution at the electrode-skin interface. We therefore compared the electric fields in the brain and the goal function values for the three different approximation approaches to reference simulations that added the electrodes as separate volumes to the head model (Fig. S3). As expected, node-wise corrections consistently resulted in very high correlations between the estimated fields and goal function values and their reference values. Interestingly, the correlations remained high (>0.98; Tables 2-5) also for the computationally simplest case of no corrections, except for large TES electrodes. In line with this, the optimizations performed similarly well independent of the chosen approximation approach, with exception of focality optimizations of large TES electrodes. Generally, the optimizations strongly outperformed standard electrode positioning and random sampling in all cases. This indicates that the optimization can be performed without current corrections for most applications (4×1 TES, TIS, ECT & TTFields) to increase computational efficiency. However, for 4×1 TES and TTFields, where several electrodes are connected to the same channel, we suggest using an electrode-wise approach in cases where the volume conduction properties of the head model might vary more strongly across electrode positions, e.g. in patients with cranial openings.

The tested application scenarios represent challenging non-convex optimization problems. Our results confirm that the chosen stochastic optimization approach, based on the differential evolution algorithm (Storn and Price, 1997), reliably achieves good results. However, its convergence generally depends on the complexity of the optimized function, so that the final goal function values for focality optimization for TIS and to a lower extent also for TTFields optimization exhibited a noticeable spread. In practice, these applications thus benefit from a multi-start approach.

### Limitations and future steps

The convergence and computational efficiency of our approach depends on the number of free parameters (up to 12 were tested here) and the complexity of the goal function. These properties together with the suitable current correction method need to be confirmed in pilot tests for new applications. In case of a high number of available stimulation channels, existing multi-electrode optimization approaches likely perform better, even though most of them are limited to the use of less complex goal functions (Dmochowski et al., 2011; Fernandez-Corazza et al., 2020; Saturnino et al. 2019).

Future work could aim at increasing the computational efficiency of the new optimization framework. Node-wise current corrections were most accurate, but were computationally also least efficient. They could be strongly improved using surrogate models to predict the node currents in dependence of the electrode positions and orientations, as already done for the electrode-wise current corrections (Suppl. Material S1). Possibly more efficient optimization algorithms could be tested, depending on the application. For example, it seems likely that the goal function for intensity optimization of focal 4×1 TES changes smoothly when varying the input parameters, so that faster local search algorithms might give similarly good results. The efficiency of the optimizations could likely be increased by directly specifying constraints of the feasible position and orientation parameters instead of penalizing the objective function. While this is challenging due to the irregularly shaped boundary of the valid head region, this might particularly help to speed up the optimization of large electrode arrays, as for TTFields.

The optimization framework could be extended to further applications and goal functions in the future. The examples of the geometry optimization of TES electrodes and the optimization of the patient-specific optimization of ECT montages were included to showcase the flexibility of our approach, but would likely need further adaptations to the specific clinical applications. Along similar lines, the efficacy of TTFields seems to depend on the orthogonality of the electric fields applied by the two electrode array pairs within the tumor (Korshoej & Thielscher, 2018; Korshoej et al., 2019a). In in vivo experiments, two sequential and orthogonal fields increased the therapeutic efficacy by ≈20% compared to one constantly active field (Kirson et al., 2007). Optimizing not only field intensity, but also the angle between the field vectors induced in the tumor by the two electrode pairs might therefore be a valuable extension of the TTFields optimization approach established here, which could be added in the future by modifying the goal function accordingly.

## 5. Code availability

The presented method is implemented in SimNIBS v4.5. We provide easy-to-use application examples for each of the investigated cases, i.e. standard TES, focal 4×1 TES, TIS and TTFields on the SimNIBS homepage (https://simnibs.github.io/simnibs/build/html/index.html). Using the *ernie* headmodel from the SimNIBS example dataset, the optimizations can be carried out on a standard notebook with 16 GB RAM.

## Acknowledgements

The project was supported by the Danish Cancer Society (R322-A17630-B5570). AT was supported by the Lundbeck Foundation (grants R313-2019-622 and R244-2017-196) and the National Institute of Health (grant R01MH128422).

## 6. Credit Authorship Statement

KW, AT, and AK formulated the overarching research goals and aims. KW developed the optimization approach and implemented it in SimNIBS together with AT and TW. KHM and KW worked on the selection and hyperparameter optimization of the optimization algorithm. AK contributed with important clinical aspects in the definition of optimization goals. KW wrote the initial draft and prepared the visualizations. AK acquired the financial support. KW and TRK provided the computing resources for the computationally expensive calculations. TRK contributed defining the intensity-focality tradeoff. All authors critically reviewed the whole manuscript.

## References

Ballo, M. T., Urman, N., Lavy-Shahaf, G., Grewal, J., Bomzon, Z. E., & Toms, S. (2019). Correlation of tumor treating fields dosimetry to survival outcomes in newly diagnosed glioblastoma: a large-scale numerical simulation-based analysis of data from the phase 3 EF-14 randomized trial. International Journal of Radiation Oncology* Biology* Physics, 104(5), 1106-1113.

Bestmann, S., & Walsh, V. (2017). Transcranial electrical stimulation. Current Biology, 27(23), R1258-R1262.

Cao, F., Madsen, K. H., Puonti, O., Siebner, H. R., Schmitgen, A., Kunz, P., & Thielscher, A. (2024) FEM-based Electric Field Calculations for Neuronavigated Transcranial Magnetic Stimulation. Biorxiv, 10.1101/2024.12.13.628139.

Chakrabarti, S., Grover, S., & Rajagopal, R. (2010). Electroconvulsive therapy: a review of knowledge, experience and attitudes of patients concerning the treatment. The World Journal of Biological Psychiatry, 11(3), 525-537.

David C. Van Essen, Stephen M. Smith, Deanna M. Barch, Timothy E.J. Behrens, Essa Yacoub, Kamil Ugurbil, for the WU-Minn HCP Consortium. (2013). The WU-Minn Human Connectome Project: An overview. NeuroImage 80(2013):62-79.

Dmochowski, J., Datta, A., Bikson, M., Su, Y., & Parra, L. (2011). Optimized multi-electrode stimulation increases focality and intensity at target. Journal of Neural Engineering, 8, 46011.

Edwards, D., Cortes, M., Datta, A., Minhas, P., Wassermann, E. M., & Bikson, M. (2013). Physiological and modeling evidence for focal transcranial electrical brain stimulation in humans: a basis for high-definition tDCS. Neuroimage, 74, 266-275.

Fernandez-Corazza, M., Turovets, S., & Muravchik, C. H. (2020). Unification of optimal targeting methods in transcranial electrical stimulation. Neuroimage, 209, 116403.

Gabriel, C., Peyman, A., & Grant, E. H. (2009). Electrical conductivity of tissue at frequencies below 1 MHz. Physics in medicine & biology, 54(16), 4863.

Grossman, N., Bono, D., Dedic, N., Kodandaramaiah, S. B., Rudenko, A., Suk, H. J.,…& Boyden, E. S. (2017). Noninvasive deep brain stimulation via temporally interfering electric fields. cell, 169(6), 1029-1041.

Kirson, E. D., Dbalý, V., Tovaryš, F., Vymazal, J., Soustiel, J. F., Itzhaki, A.,…& Palti, Y. (2007). Alternating electric fields arrest cell proliferation in animal tumor models and human brain tumors. Proceedings of the National Academy of Sciences, 104(24), 10152-10157.

Korshoej, A. R., Hansen, F. L., Thielscher, A., von Oettingen, G. B., & Sørensen, J. C. H. (2017). Impact of tumor position, conductivity distribution and tissue homogeneity on the distribution of tumor treating fields in a human brain: A computer modeling study. PloS one, 12(6), e0179214.

Korshoej, A. R., Hansen, F. L., Mikic, N., von Oettingen, G., Sørensen, J. C. H., & Thielscher, A. (2018). Importance of electrode position for the distribution of tumor treating fields (TTFields) in a human brain. Identification of effective layouts through systematic analysis of array positions for multiple tumor locations. PLoS One, 13(8), e0201957.

Korshoej, A. R., & Thielscher, A. (2018). Estimating the intensity and anisotropy of tumor treating fields using singular value decomposition. Towards a more comprehensive estimation of anti-tumor efficacy. In 2018 40th Annual international conference of the IEEE Engineering in Medicine and Biology Society (EMBC) (pp. 4897-4900). IEEE.

Korshoej, A. R., Sørensen, J. C. H., Von Oettingen, G., Poulsen, F. R., & Thielscher, A. (2019a). Optimization of tumor treating fields using singular value decomposition and minimization of field anisotropy. Physics in Medicine & Biology, 64(4), 04NT03.

Korshoej, A. R., Mikic, N., Hansen, F. L., Saturnino, G. B., Thielscher, A., & Bomzon, Z. E. (2019b). Enhancing tumor treating fields therapy with skull-remodeling surgery. The role of finite element methods in surgery planning. In 2019 41st annual international conference of the IEEE Engineering in Medicine and Biology Society (EMBC) (pp. 6995-6997). IEEE.

Korshoej, A. R., Lukacova, S., Lassen-Ramshad, Y., Rahbek, C., Severinsen, K. E., Guldberg, T. L.,…& Sørensen, J. C. H. (2020). OptimalTTF-1: Enhancing tumor treating fields therapy with skull remodeling surgery. A clinical phase I trial in adult recurrent glioblastoma. Neuro-oncology advances, 2(1), vdaa121.

Martin, D. M., Bakir, A. A., Lin, F., Francis-Taylor, R., Alduraywish, A., Bai, S.,…& Loo, C. K. (2021). Effects of modifying the electrode placement and pulse width on cognitive side effects with unilateral ECT: A pilot randomised controlled study with computational modelling. Brain Stimulation, 14(6), 1489-1497.

Mikic, N., Poulsen, F. R., Kristoffersen, K. B., Laursen, R. J., Guldberg, T. L., Skjøth-Rasmussen, J.,…& Korshøj, A. R. (2021). Study protocol for OptimalTTF-2: enhancing Tumor Treating Fields with skull remodeling surgery for first recurrence glioblastoma: a phase 2, multi-center, randomized, prospective, interventional trial. BMC cancer, 21(1), 1-8.

Mikic, N., Gentilal, N., Cao, F., Lok, E., Wong, E. T., Ballo, M.,…& Korshoej, A. R. (2024). Tumor-treating fields dosimetry in glioblastoma: Insights into treatment planning, optimization, and dose–response relationships. Neuro-Oncology Advances, 6(1), vdae032.

Miranda, P. C., Lomarev, M., & Hallett, M. (2006). Modeling the current distribution during transcranial direct current stimulation. Clinical Neurophysiology, 117(7), 1623–1629. 10.1016/j.clinph.2006.04.009

Mun, E. J., Babiker, H. M., Weinberg, U., Kirson, E. D., & Von Hoff, D. D. (2018). Tumor-treating fields: a fourth modality in cancer treatment. Clinical Cancer Research, 24(2), 266-275.

Opitz, A., Paulus, W., Will, S., Antunes, A., & Thielscher, A. (2015). Determinants of the electric field during transcranial direct current stimulation. Neuroimage, 109, 140-150.

Panou, G., & Korakitis, R. (2019). Geodesic equations and their numerical solution in Cartesian coordinates on a triaxial ellipsoid. Journal of Geodetic Science, 9(1), 1-12.

Puonti, O., Van Leemput, K., Saturnino, G. B., Siebner, H. R., Madsen, K. H., & Thielscher, A. (2020). Accurate and robust whole-head segmentation from magnetic resonance images for individualized head modeling. Neuroimage, 219, 117044.

Saturnino, G. B., Siebner, H. R., Thielscher, A., & Madsen, K. H. (2019). Accessibility of cortical regions to focal TES: Dependence on spatial position, safety, and practical constraints. NeuroImage, 203, 116183.

Saturnino, G. B., Madsen, K. H., & Thielscher, A. (2019a). Electric field simulations for transcranial brain stimulation using FEM: an efficient implementation and error analysis. Journal of neural engineering, 16(6), 066032.

Saturnino, G. B., Madsen, K. H., & Thielscher, A. (2021). Optimizing the electric field strength in multiple targets for multichannel transcranial electric stimulation. Journal of neural engineering, 18(1), 014001.

Schenk, O., Gärtner, K. (2011). PARDISO. In: Padua, D. (eds) Encyclopedia of Parallel Computing. Springer, Boston, MA. 10.1007/978-0-387-09766-4_90

Segar, D. J., Bernstock, J. D., Arnaout, O., Bi, W. L., Friedman, G. K., Langer, R.,…& Rampersad, S. M. (2023). Modeling of intracranial tumor treating fields for the treatment of complex high-grade gliomas. Scientific Reports, 13(1), 1636.

Souza, V. H., Matsuda, R. H., Peres, A. S., Amorim, P. H. J., Moraes, T. F., Silva, J. V. L., & Baffa, O. (2018). Development and characterization of the InVesalius Navigator software for navigated transcranial magnetic stimulation. Journal of neuroscience methods, 309, 109-120.

Storn, R., & Price, K. (1997). Differential evolution–a simple and efficient heuristic for global optimization over continuous spaces. Journal of global optimization, 11, 341-359.

Virtanen, P., Gommers, R., Oliphant, T. E., Haberland, M., Reddy, T., Cournapeau, D.,…& Van Mulbregt, P. (2020). SciPy 1.0: fundamental algorithms for scientific computing in Python. Nature methods, 17(3), 261-272.

Wagner, T. A., Zahn, M., Grodzinsky, A. J., & Pascual-Leone, A. (2004). Three-dimensional head model simulation of transcranial magnetic stimulation. IEEE Transactions on Biomedical Engineering, 51(9), 1586-1598.

Weise, K., Poßner, L., Müller, E., Gast, R., & Knösche, T. R. (2020). Pygpc: a sensitivity and uncertainty analysis toolbox for Python. SoftwareX, 11, 100450.

Wenger, C., Miranda, P. C., Salvador, R., Thielscher, A., Bomzon, Z., Giladi, M.,…& Korshoej, A. R. (2018). A review on tumor-treating fields (TTFields): clinical implications inferred from computational modeling. IEEE Reviews in Biomedical Engineering, 11, 195-207.

Zienkiewicz, O. C., & Zhu, J. Z. (1992). The superconvergent patch recovery and a posteriori error estimates. Part2: Error estimates and adaptivity. International Journal for Numerical Methods in Engineering, 33(7), 1365-1382.

