## Supplemental Material for "A Leadfield-Free Optimization Framework for Transcranially Applied Electric Currents"

Konstantin Weise[1,2,3*], Kristoffer H. Madsen[4,5], Torge Worbs[5,7], Thomas R. Knösche[3], Anders Korshøj[1,6#], Axel Thielscher[5,7#]

[1]Department of Clinical Medicine, Aarhus University, Aarhus, Denmark

[2]Methods and Development Group "Brain Networks", Max Planck Institute for Human Cognitive and Brain Sciences, Leipzig, Germany.

[3]Leipzig University of Applied Sciences (HTWK), Institute for Electrical Power Engineering, Leipzig, Germany

[4]Technical University of Denmark, Section for Cognitive Systems, Department of Applied Mathematics and Computer Science, Kongens Lyngby, Denmark.

[5]Danish Research Centre for Magnetic Resonance, Department of Radiology and Nuclear Medicine, Copenhagen University Hospital Amager and Hvidovre, Hvidovre, Denmark.

[6]Department of Neurosurgery, Aarhus University Hospital, Aarhus, Denmark

[7]Technical University of Denmark, Section for Magnetic Resonance, Department of Health Technology, Kongens Lyngby, Denmark.

*CORRESPONDING AUTHOR

[#]contributed equally

### S1. Modeling of Interface Currents

Setting von Neumann boundary conditions at the skin nodes to approximate the current flow at the electrode-skin interfaces avoids the need for repeating the FEM preparation steps and decreases the costs for evaluating the goal function for new parameter choices. However, it requires setting the node currents $I_1$, $I_2$, *etc.* (Fig. 1 of the main manuscript) at the interface areas correctly. In particular, applying a constant current density does not correctly represent the underlying electric field problem for TES with large electrodes, in which the currents injected into an electrode cause an approximately constant electric potential (rather than constant current density) at the skin interface. When several electrodes are connected to the same stimulation channel (as in case of TTFields), then the skin interfaces of all those electrodes will share a common electric potential. While modeling a constant electric potential at one or several interfaces is easily feasible by applying Dirichlet boundary conditions to define the potentials of the corresponding nodes ($V_1$, $V_2$ in Fig. 1), this requires updates of the stiffness matrix and repetition of the preparation steps of the solver, thus compromising efficiency.

*Node-wise Dirichlet approximation:* In order to avoid the need for explicitly setting Dirichlet boundary conditions, we instead solve a secondary optimization problem to iteratively tune the node currents $I_1$, $I_2$, … when updating the electrode positions or shapes so that the conditions for constant electric potentials at the skin interfaces of electrodes sharing the same stimulation channel are met. This approach is referred to as *node-wise Dirichlet approximation* in the following and, while requiring repeated FEM solutions, is much more efficient than setting Dirichlet boundary conditions directly. In this case, the secondary optimization problem must be solved for all involved skin nodes to achieve the correct, inhomogeneous current density distribution at the electrode-skin interfaces, as can be seen in Fig. 1.

*Electrode-wise Dirichlet approximation*: The impact of the inhomogeneous current distribution at the electrode interfaces on the resulting electric field in the brain depends on the size and relative position of the electrodes to each other and generally is much weaker for small electrodes used for TTF and focal 4x1 TES than for the large electrodes in standard unfocal TES experiments (e.g. with edge lengths of 50-70 mm). Neglecting this inhomogeneity represents a simplification of the optimization problem and allows for a significant acceleration of its solution. If it is assumed that the current density is constant over the entire electrode, then in cases were several electrodes share a common channel, only the total currents flowing into the electrode need to be optimized instead of the many individual currents of the single nodes. Using the example in Fig. 1, this would be the four outer currents $I_1 \ldots I_4$. This approach is referred to as *electrode-wise Dirichlet approximation*. We have investigated the extent to which this approach is applicable and how it affects the computing time and the accuracy of the electric field in the brain for various applications and electrode sizes.

The correction function of the electrode-wise currents depends on the position and orientation of the electrodes and is expected to be continuous and differentiable over the skin surface. The information about this correction function can be monitored and used to continuously improve the starting conditions during the entire optimization process, thereby significantly reducing the number of iterations required to solve the current optimization problem. Therefore, we construct surrogate models, which learn from optimized currents during the whole optimization process to predict the currents of the next position and orientation of the electrode array. For this purpose, we used the generalized polynomial chaos expansion (gPC) method implemented in the Python package *pygpc* (Weise et al., 2020). The number of Dirichlet correction iterations is shown as an example in Fig. S1 for the case of a TTF array with 3x3 electrodes. If the model is untrained, between 8 and 14 additional calculations are initially required to determine the correct currents to ensure a uniform voltage across all electrodes. As the optimization progresses, more and more training data becomes available, which can even reduce the number to such an extent that in some cases no additional calculations need to be performed.

Currently, the method can only be applied to electrode-wise current correction, as the number of free parameters (number of electrodes) does not change depending on the electrode position. With node-wise correction, on the other hand, the number of nodes assigned to the electrodes differ for each electrode position due to the discretization of the skin surface. This means that a clear assignment of the nodes across the parameter space is no longer possible for each electrode position.

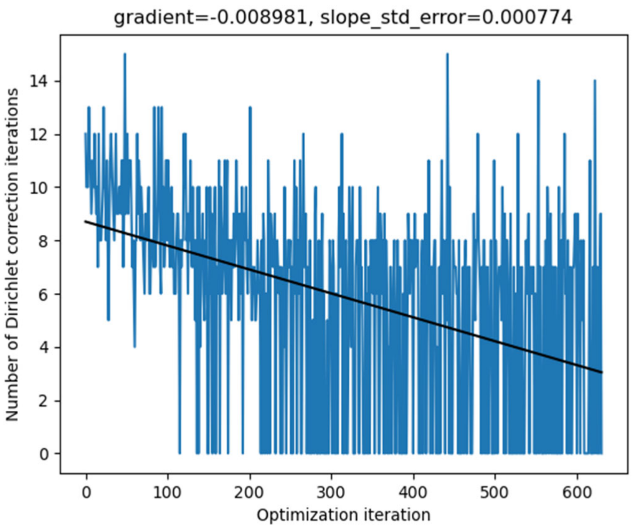


**Fig. S1:** Number of Dirichlet correction iterations to determine the currents per electrode to ensure the same voltage for a TTF array containing 3x3 electrodes over the course of an optimization run.

### S2. Ellipsoidal Coordinate System of the Head

The optimization of the electrode arrangement requires a suitable parameterization and automated assignment of the corresponding skin surface nodes to the associated electrodes by the optimization algorithm. However, the electrodes should be placed at certain positions of the skin surface, such as the face or the ears. For this reason, an individual mask from an MNI template is created during the initialization of the optimizer, which automatically selects the area for valid electrode positions. Electrode configurations that are outside this area, partially overlap with the border, or overlap among themselves will not be considered and penalized during the optimization.

The geometrical arrangement of the electrodes in the array is defined by the user in a normalized 2D space ($xy$-plane, Fig. 2 of the main manuscript). In order to be able to move the array over the head surface in a parameterized manner, a triaxial ellipsoid is fitted to the nodes of the valid skin region in a least square sense. The center of the electrode array is then given by the spherical coordinates $(\theta', \varphi')$ shown in Fig. 2. The orientation of the array is defined by the angle $\alpha$ between the electrode reference direction ($y$-axis in normalized electrode space) and the vector of constant $\varphi$ in the ellipsoidal space. The task is to project the electrode array from the normalized $xy$-plane onto the triaxial ellipsoid and to identify the corresponding skin nodes subsequently. Given the position and orientation of the electrode array on the triaxial ellipsoid, the position of the electrodes can be determined by solving the associated direct geodesic problem, known from the field of differential geometry, as depicted in Fig. 2. This results in solving the geodesic initial value problem on a triaxial ellipsoid given the starting point from the electrode center $(\theta', \varphi')$ into the direction $\vec{s}$ for a given distance $d$ to the center of the target electrode $(\theta'', \varphi'')$ whose position is to be determined. This has to be done for all electrodes in the array in every iteration. The problem was solved by using the approach of Panou and Korakitis (2019). The computation time is comparatively short, with around 0.05 sec for a 4x1 TES montage with 4 external electrodes. Once the positions of the electrodes on the ellipsoid are chosen for a particular iteration during optimization, the corresponding positions on the head surface are determined. Rays are directed from the ellipsoid in the normal direction towards the head surface and the exact points of intersection with the skin surface are determined. Matching masks in the form of the user electrodes are then placed around the intersecting points and the enclosed nodes and triangles are determined.

### S3. Optimization approach

Once the optimization problems and their constraints are defined, a suitable optimization algorithm must be selected to solve the problems. Because the electrode arrays are free to rotate it is expected that the objective functions are not convex and have numerous local minima. For this reason, a stochastic optimization approach based on the differential evolution algorithm (Storn and Price, 1997) was chosen. Here, we used the `differential_evolution` algorithm implemented in *SciPy* (Virtanen et al., 2020).

The convergence properties of the algorithm depend on the hyperparameters *mutation*, *recombination* and *population size*. By means of numerous test simulations, the following parameters were identified that provided good and reliable convergence across all problem classes. The mutation parameter introduces diversity into the sampled population and modifies the parameters of the mutants randomly. It was chosen to dither randomly between (0.01, 0.5) according to a uniform distribution on a generation-by-generation basis. The recombination parameter controls the number of mutants to progress into the next generation and was set to 0.7. Finally, the population size defines the number of mutants in the population and was set to 13.

The procedure for the entire optimization process is outlined in the following:

Preparation phase (initialization):

- ROI definition: The ROIs (and non-ROIs) are defined based on user defined points, surfaces, or volumes in the head model (structured or scattered points). For each ROI, a matrix $[\mathbf{S}]$ for fast interpolation of the electric field is pre-calculated and stored.
- Prepare FEM: The stiffness matrix is computed for isotropic or anisotropic conductivity profiles, preconditioned, and stored.
- Determine valid skin region where electrodes are allowed to be placed

- Fit the ellipsoid
- Create an initial random population representing different electrode configurations

The steps in each optimization iteration are:

- Create a new random population, based on the previous with several mutants representing different electrode configurations, each with parameter set $\boldsymbol{x}$ to test.
- Determine the associated skin nodes depending on the electrode locations to inject the source currents.
- Set source terms on the right-hand side (RHS) of the system of equations by assigning currents to the selected nodes according to their equivalent areas and weights.
- Solve FEM: Determine the electric potential in the nodes by solving the system of equations using the PARDISO solver (Schenk und Gärtner, 2011) of the MKL library (http://software.intel.com/en-us/intel-mkl).
- Apply Dirichlet correction: Check if node voltages over electrodes of the same channel are equal, otherwise correct the node currents and solve again. If voltages over electrodes do not vary within a tolerance interval of 1%, continue, otherwise refine the currents and check voltages again.
- Determine electric field in all tetrahedral elements from the node potentials by computing its gradient over the tetrahedral elements.
- Interpolate electric field to ROIs by applying the precomputed interpolation matrices.
- Compute the QOI of the electric field in the ROIs, e.g. magnitude, normal, or tangential component.

- Determine the (application specific) goal function value and pass it to the optimizer.

The process is repeated until the objective functions no longer change within a specified relative tolerance interval of 0.1 and no further substantial improvement can be observed. At the end, a standard SimNIBS simulation with full modeling of the electrodes is run to determine the final electric field distribution for the optimized parameters.

### S4. Verification of the triaxial ellipsoid fitting procedure

The parameterization of the electrode arrays is based on the geodesic coordinates resulting from fitting a triaxial ellipsoid to the part of the skin surface, where electrodes can be applied.

The stability of the fitting procedure is investigated using head shapes with extraordinary geometrical extent in anterior-poster, lateral-medial, and superior-inferior direction, which maximize the ratio between the longest and the shortest semi-axis. The head models were chosen from the connectome dataset (Van Essen et al. 2013) and triaxial ellipsoids were fitted to the skin surface, where electrodes can be applied. The qualities of the fits are evaluated by applying an affine transformation of the skin points to the unit sphere using the respective ellipsoid parameters. Subsequently, the normalized mean radius (truth=1) and the standard deviation of the skin points to the transformed unit sphere were calculated. The results are shown in Fig. S2.

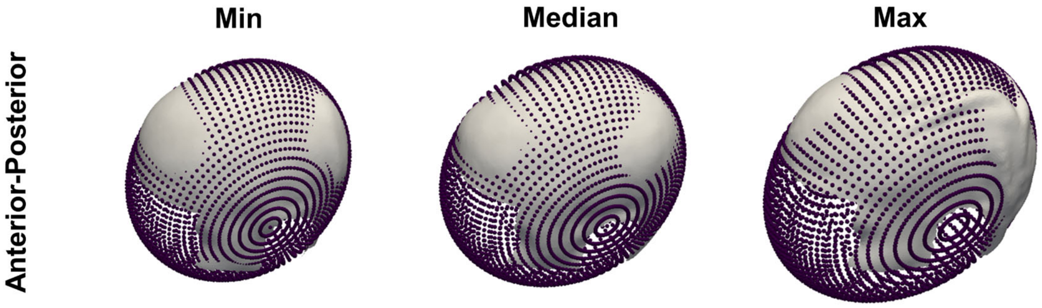


| | | | |
|---|---|---|---|
| Mean($\tilde{r}$) | 0.99 | 1.00 | 1.04 |
| Std($\tilde{r}$) | 0.07 | 0.11 | 0.20 |

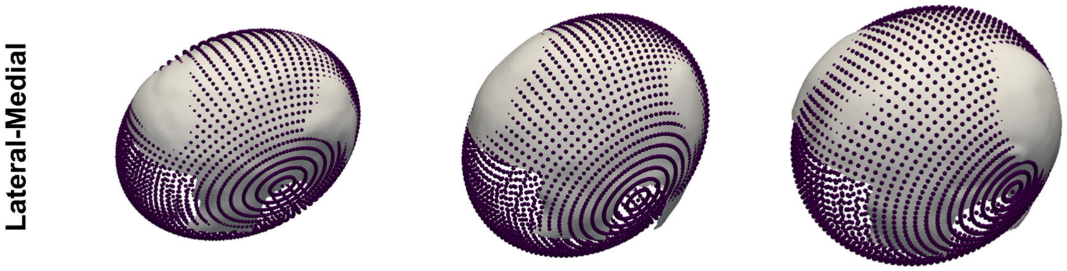


| | | | |
|---|---|---|---|
| Mean($\tilde{r}$) | 1.03 | 1.00 | 1.00 |
| Std($\tilde{r}$) | 0.17 | 0.10 | 0.08 |

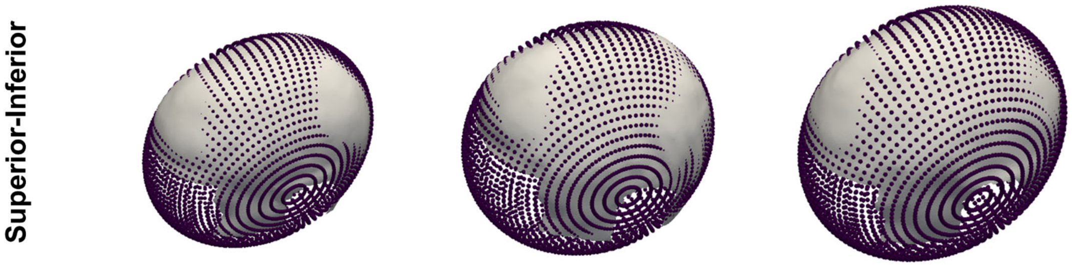


| | | | |
|---|---|---|---|
| Mean($\tilde{r}$) | 1.01 | 1.01 | 1.01 |
| Std($\tilde{r}$) | 0.13 | 0.12 | 0.11 |

**Fig. S2:** Results of fitting triaxial ellipsoids to head shapes with extraordinary extents in anterior-poster, lateral-medial, and superior-inferior direction. The normalized mean radius (truth=1) and the standard deviation of the skin points to the transformed unit sphere is given below each case.

### S5. Influence of Dirichlet correction approach on the electric field accuracy

For the various applications, the electric field in gray and white matter is compared between different Dirichlet correction approaches and reference simulations where the electrodes were modeled as separate volumes. For each case, the electric field from 200 random electrode positions was computed. The differences between the approaches are quantified using the normalized root mean square deviation (NRMSD):

$$NRMSD = \frac{\sqrt{\frac{1}{N}\sum_{i=1}^{N}\left(E_i - E_{i,ref}\right)^2}}{\left(E_{ref}\right) - \left(E_{ref}\right)}$$

where $N$ is the total number of tetrahedra in gray and white matter, $E_i$ denotes the electric field magnitude from the respective approach in the $i$-th tetrahedra, and $E_{ref}$ is the electric field of the reference solution. The NRMSD is computed for each of the 200 electrode positions and the resulting distributions are shown in Fig. S3.

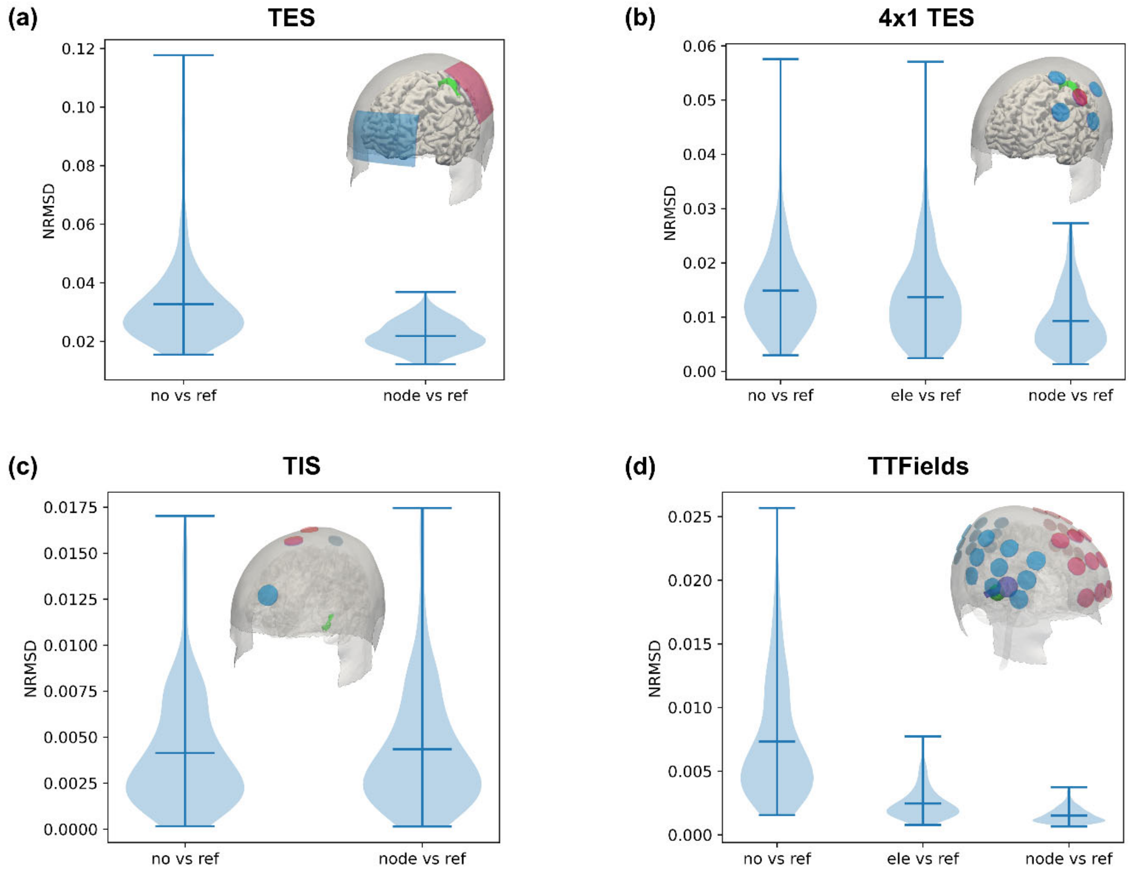


**Fig. S3:** Comparison of the electric field between different Dirichlet correction methods and reference simulations for (a) standard TES, (b) 4x1 TES, (c) TIS, and (d) TTFields.

### References


David C. Van Essen, Stephen M. Smith, Deanna M. Barch, Timothy E.J. Behrens, Essa Yacoub, Kamil Ugurbil, for the WU-Minn HCP Consortium. (2013). The WU-Minn Human Connectome Project: An overview. NeuroImage 80(2013):62-79.

Panou, G., & Korakitis, R. (2019). Geodesic equations and their numerical solution in Cartesian coordinates on a triaxial ellipsoid. Journal of Geodetic Science, 9(1), 1-12.

Schenk, O., Gärtner, K. (2011). PARDISO. In: Padua, D. (eds) Encyclopedia of Parallel Computing. Springer, Boston, MA. https://doi.org/10.1007/978-0-387-09766-4_90

Storn, R., & Price, K. (1997). Differential evolution–a simple and efficient heuristic for global optimization over continuous spaces. Journal of global optimization, 11, 341-359.

Van Essen, D. C., Smith, S. M., Barch, D. M., Behrens, T. E., Yacoub, E., Ugurbil, K., & Wu-Minn HCP Consortium. (2013). The WU-Minn human connectome project: an overview. *Neuroimage*, *80*, 62-79.

Virtanen, P., Gommers, R., Oliphant, T. E., Haberland, M., Reddy, T., Cournapeau, D., ... & Van Mulbregt, P. (2020). SciPy 1.0: fundamental algorithms for scientific computing in Python. Nature methods, 17(3), 261-272.

Weise, K., Poßner, L., Müller, E., Gast, R., & Knösche, T. R. (2020). Pygpc: a sensitivity and uncertainty analysis toolbox for Python. SoftwareX, 11, 100450.
